# Receptor stoichiometry predicts artery-typical vulnerability to altered Notch signaling during smooth muscle differentiation

**DOI:** 10.1101/2025.05.05.652186

**Authors:** Sami Sanlidag, Noora Virtanen, Hesam Hoursan, Tommaso Ristori, Marika Sjöqvist, Sandra Loerakker, Cecilia Sahlgren

## Abstract

The development and maintenance of arterial smooth muscle cells (SMCs) rely on Jagged1-Notch2/Notch3 signaling. While Notch2 and Notch3 are thought to function redundantly during SMC development, clinical and experimental evidence suggests artery-specific importance for the two receptors. Combining *in vitro*, *in vivo,* and *in silico* models, we report that the canonical Notch signaling during SMC differentiation is largely driven by Notch2. While Notch2 and Notch3 co-regulate a large group of genes in SMCs upon Jagged1 interaction, Notch3 is a less potent inducer of Notch signaling than Notch2 and requires higher doses to potentiate a meaningful transcriptional response due to its weak interaction with RBPJκ. Consequently, Notch2 depletion abolishes Notch signaling in the SMCs of the large elastic arteries. However, high Jagged1 and Notch3 expression in smaller arteries like those in the brain can compensate for Notch2 loss. This work refines our mechanistic understanding of Notch signaling in the SMCs and offers region-specific insights into the Notch-related arterial diseases.

## INTRODUCTION

Vascular smooth muscle cells (SMCs) form a multilayered wall around the endothelium in larger caliber blood vessels, such as arteries, arterioles, and veins, ensuring vascular integrity and function. The dysregulation of SMCs is linked to severe vascular complications and constitutes high clinical significance. However, our understanding of the mechanisms regulating the SMC differentiation in different arteries is incomplete, hindering the development of appropriate therapies for SMC-related pathologies.

During development, vascular SMCs proliferate and produce the extracellular matrix around the endothelium(1, 2), followed by SMC maturation. The mature SMCs are contractile and contribute to the vascular tone. In case of alterations in the vascular environment, they can switch between varying phenotypes to partake in vascular adaptation through several mechanisms, including proliferation(3). Notch signaling, a contact-dependent cell-cell signaling pathway, is a master regulator of SMC development and homeostasis(4). In mammals, Notch signaling comprises four transmembrane Notch receptors (Notch1, 2, 3, and 4) and five ligands (Jagged1 and 2; and Delta-like 1,3, and 4). When a ligand pulls on a Notch receptor, the receptor becomes poised for two consequent cleavages mediated by ADAM (S2 cleavage) and γ-secretase (S3 cleavage), respectively. This liberates the active Notch intracellular domain (NICD) that can translocate to the nucleus and drive transcription through its co-factors, such as RBPJκ(5, 6). Generally, the receptor-ligand interactions between two juxtaposed cells (trans-interactions) activate the receptor, while those on the same cell (cis-interactions) are considered inhibitory(7). Notch signaling drives the specification and expansion of SMCs by regulating platelet-derived growth factor receptor β (Pdgfrβ) levels(8, 9) and controls their maturation by regulating contractile gene expression and quiescence(10, 11). Furthermore, it propagates around the endothelium through a mechanism called lateral induction, where the Notch activity driven by the endothelial Jagged1 leads to the expression of the SMC-intrinsic Jagged1, which activates Notch in the surrounding cells, thereby contributing to the vascular wall assembly(12–14).

Vascular SMCs mainly express receptors Notch2 and Notch3 and ligand Jagged1. Dysregulation of these molecules can cause vascular complications affecting different vessels in the arterial tree. Alagille syndrome (ALGS) is a congenital disorder attributed to mutations in Jagged1 and, less frequently, Notch2(15). It manifests with multisystem complications all around the body, including intracranial hemorrhages, aneurysms, and outflow tract defects (16–18). Notch3 is linked to perivascular complications in pulmonary hypertension, such as hypermuscularization and stenosis(19). Cerebral autosomal dominant arteriopathy with subcortical infarcts and leukoencephalopathy (CADASIL) is caused by mutations in Notch3 and manifests with dementia and smooth muscle loss in smaller caliber arteries and arterioles(20, 21). Similarly, transgenic animals lacking these proteins show perivascular pathology. Mice with endothelial cell-specific knockout of Jagged1 are deficient in SMCs and die early during embryonic development(22). Notch2 hypomorphic (Notch2^del1/del1^) mice show abnormalities, such as attenuated contractile marker expression and structural defects in the larger elastic pulmonary arteries and aorta, as well as significantly increased rates of patent ductus arteriosus (failure in the closure of the ductus arteriosus) and mortality shortly after birth(10, 23, 24). Notch3 knockout does not cause a lethal periarterial phenotype in the developing mouse, apart from mild reduction in the late contractile markers(25, 26) and later defects in the myogenic tone and SMC distribution in small vessels, such as those in the tail and the brain (27–29). However, the combination of Notch2 hypomorphism and Notch3 knockout is embryonically lethal due to severe perivascular defects, implying overlapping functions between the two receptors (10). There is also evidence supporting opposing roles for Notch2 and Notch3 in vascular SMC growth and homeostasis. Notch3 promotes proliferation and survival in concert with the ERK signaling(30) *in vitro,* and its elevated activity is linked to the overgrowth of SMCs during pulmonary hypertension(19) and in the aorta of elastin-null mice(31). Meanwhile, Notch2 promotes cell cycle arrest by p27 stabilization(11). Despite these indications, a systematic understanding of how Jagged1 signals through Notch2 and Notch3 to regulate different cellular processes during vascular SMC development has been missing.

In this work, we sought to understand the signaling mechanism involving Jagged1, Notch2, and Notch3 and how they regulate SMC differentiation in different arteries in the body. First, using *in vitro* assays and a whole transcriptome analysis, we found that Notch2 and Notch3 co-regulated transcriptional programs relating to periarterial development following Jagged1 induction in a time-dependent manner. However, the two receptors differed in their ability to potentiate transcription. Notch2 and Notch3 responded similarly to Jagged1 at the protein level, but Notch2 ICD was transcriptionally more robust. Notch3 ICD could reach substantial transcriptional activity only when it was present at high levels. Furthermore, through *in situ* hybridization and single-cell transcriptomic databases, we found that SMCs at different arteries varied highly in Notch3 and Jagged1 expression. By integrating these findings into a computational model of Notch signaling in arteries, we predicted artery-typical vulnerability to altered Notch signaling in the SMCs, where hypoactivity of Notch2 and Jagged1, but not of Notch3, severely affected Notch signaling. However, SMCs of the arteries with high levels of Notch3 and Jagged1 were predicted to be less dependent on Notch2. Contractile gene reporter assays in zebrafish larvae injected with Notch2 and Notch3 morpholinos recapitulated these predictions, showing that Notch3 can compensate for Notch2 depletion in SMCs of the cerebral vessels that express high levels of Notch3. Overall, our work provides clues for how the altered Jagged1-Notch2 and -Notch3 axes can contribute to human periarterial disease onset with local specificity.

## RESULTS

### Notch2 and Notch3 act as functional analogs during Jagged1-mediated vascular smooth muscle differentiation, but Notch2 is more robust

To systematically address the functions of Notch2 and Notch3 in the Jagged1-mediated SMC phenotypic regulation, we chose an *in vitro* approach and used immobilized recombinant human Jagged1-Fc dimeric proteins (J1-Fc) to activate Notch signaling in human aortic SMCs, followed by bulk RNA sequencing (Fig. 1A). First, we silenced Notch2 or Notch3 in the SMCs using siRNAs (siN2, siN3). 48h post-transfection, Notch2 and Notch3 protein levels were significantly reduced compared to the cells transfected with the non-targeting control (siC) (Fig. 1B), demonstrating successful silencing of the receptors. We saw a slight reduction in the Notch3 levels with siN2 (Fig. 1B; S1A,K). This was expected since Notch3 is a transcriptional target of Notch2(10). Neither of the siRNAs led to an appreciable change in Notch1 mRNA (Fig. S1A). We then plated the cells on immobilized J1-Fc and harvested them for RNA extraction after 4 and 24 hours to capture an early, more Notch-specific transcriptional response, and a later response relating to the SMC differentiation program, respectively. By setting thresholds of fold change (FC) >2 and false discovery rate (FDR) <0.05, we identified 78 genes at 4h (47 up- and 31 downregulated) and 583 genes at 24h (249 up- and 334 downregulated) differentially regulated by Jagged1 induction in the control cells. The top 10 up- and downregulated genes are annotated in Fig. 1C (See Dataset S1 for a complete list of the DEGs).

**Figure 1.**
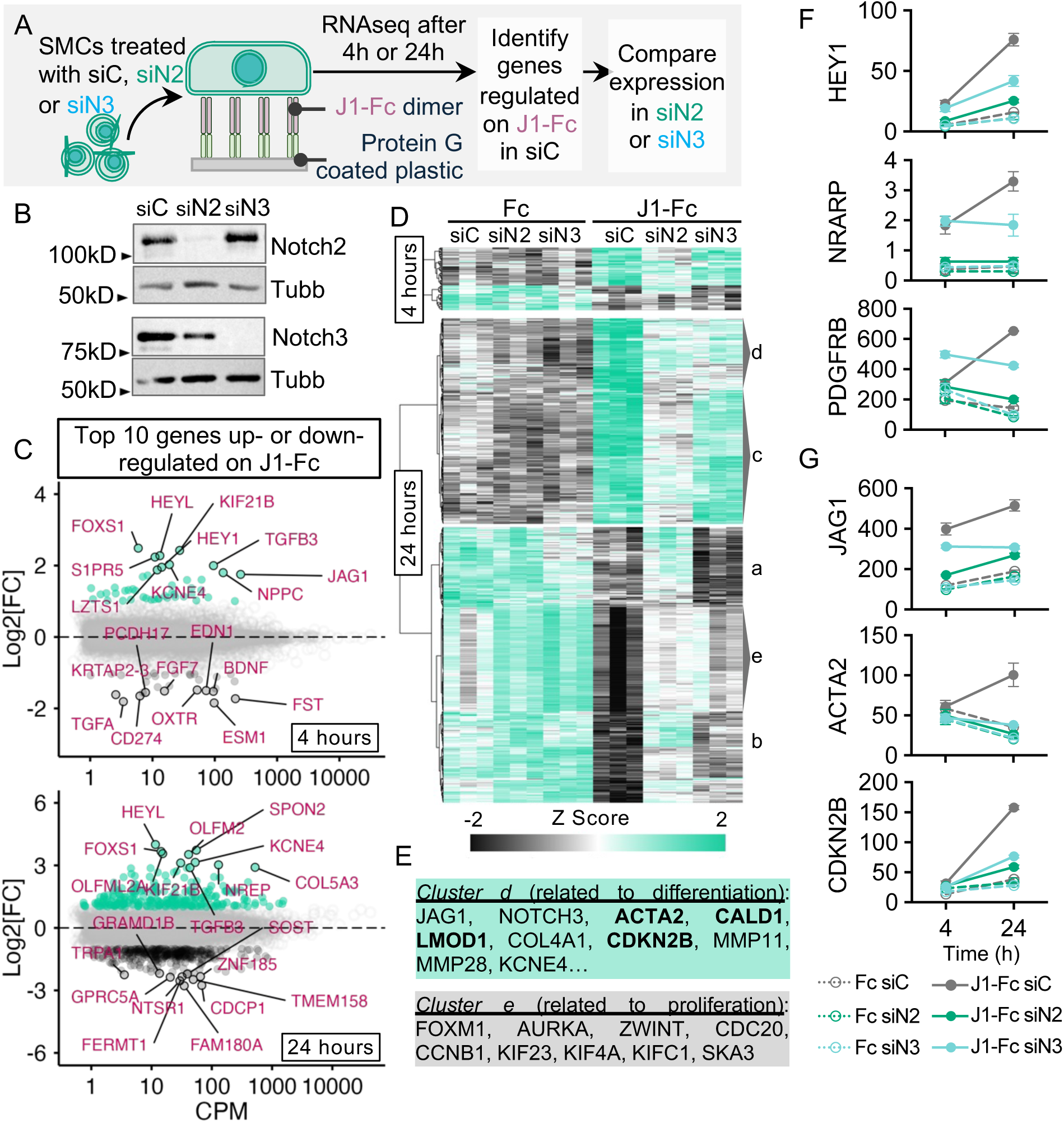
Notch2 and Notch3 act as functional analogs during Jagged1-mediated SMC differentiation, but Notch2 facilitates the early transcriptional response. (A) Schematic description of the experimental setup and the analysis strategy for the RNAseq experiment. (B) Notch2 and Notch3 protein levels in SMCs 48h post-transfection with the corresponding siRNAs. (C) Genes significantly regulated in SMCs on immobilized J1-Fc at 4h and 24h (FC>2, FDR>0.05) based on RNAseq. The top 10 up- or downregulated genes are annotated (n=3 independent experiments). (D) Heatmaps clustered by Ward’s method showing the z-scores of genes regulated in SMCs on immobilized J1-Fc at 4h and 24h time points across cells treated with control, Notch2, or Notch3 siRNAs (n=3 independent experiments). Gene clusters were considered as follows: Cluster c, upregulated more by Notch2 and less by Notch3; cluster d, upregulated by both Notch2 and Notch3; and Cluster e, downregulated by both Notch2 and Notch3. (E) Representative genes from the clusters on J1-Fc (Cluster d) and genes related to proliferation downregulated on J1-Fc (Cluster e). (F-H) Expression of genes representative of Notch signaling activity, contractile differentiation, and cell cycle arrest (mean±SD).

At 4h, many putative Notch targets, such as HEY1, HEYL, HES1, NRARP, and JAG1, as well as genes related to perivascular identity (KCNE4(32)), SMC quiescence (CDKN2B(33)), and TGFβ signaling effectors (TGFB3 and SMAD3), were upregulated. Interestingly, the silencing of Notch2 and not of Notch3 led to a dramatic loss of transcriptional response to Jagged1, as evidenced by the overall behavior of the gene cluster across the groups (Fig. 1D; S1D). This pattern was reflected in many of the Notch targets, such as HEY1 and NRARP (Fig. 1F). Only a few genes, such as KCNE4 and TGFB3, seemed to be affected by Notch3 silencing to a similar degree to Notch2 (Fig. S1A) even though these cells express NOTCH3 mRNA approximately twice as much as Notch2 under basal conditions (Fig. S3A).

At 24h, the earlier markers of SMC contractility such as ACTA2, CALD1, and LMOD1, but not the mature SMC markers, such as MYH11 and SMTN (Fig. 1 A), were among the upregulated genes, confirming an intermediate stage of contractile differentiation following the receptor activation by immobilized Jagged1. On the other hand, the transcriptional response was convoluted at this time point, and it was difficult to discern between the Notch2- and Notch3-silenced groups using simple statistical comparison. For easier interpretation, we divided the DEGs at 24h into five Ward clusters (2 up- and 3 downregulated clusters) (Fig. 1D, also see Dataset S1 for the clusters). Both upregulated clusters, *c* and *d*, were enriched for ontology terms related to vascular myogenic processes, while the three downregulated clusters, *a*, *b,* and *e,* were enriched for vascular process-, synaptic signaling-, and cell cycle-related terms, respectively (Fig. S1C).

Notch2 silencing also had a profound impact on all clusters at 24h. This effect was particularly evident in *cluster a* and the majority of *cluster c (See Fig. S1D for cluster behavior),* which were affected little by Notch3 silencing. SNAI2 (Fig. S1B), an important transcription factor for the recruitment of mural progenitors around the aorta from the sclerotome, was also specifically regulated by Notch2 silencing (*cluster c*) (34). *Cluster a* included genes such as SOST, RGS2, and EDNRB, which mark different zones in the pulmonary-aortic tree(35), and genes linked to adverse aortic remodeling, FERMT1, and BMP2 (See Dataset S1) (36, 37) - posing the question of whether Notch2 might be important for the identity of the SMCs at this location.

Notch3 silencing elicited a consistent (albeit smaller compared to Notch2) reduction in the ligand-induced transcriptional response at 24h. Overall, it seemed that many genes responded strongly to Notch2 already at 4h, while Notch3 only added to this response at 24h. This pattern was reflected in some genes of *cluster c,* including Notch targets NRARP and HEY1; the perivascular marker PDGFRB (Fig. 1F); and the extracellular matrix components, such as COL5A1 (Fig. S1B). The impact of Notch3 was most evident in c*lusters d* and *e,* which responded to the silencing of Notch2 and Notch3 similarly (Fig. 1D; S1D). These clusters were enriched for genes related to the hallmarks of SMC maturation: contractile markers, extracellular matrix deposition (e.g., COL4A1), and cell cycle arrest (e.g., CDKN2B) (Fig. 1G; S1A). Interestingly, the optimal expression of genes related to SMC contractility, such as ACTA2 encoding for α-smooth muscle actin (αSMA), seemed to rely on both receptors (Fig. 1G). Notch3 knockout mice develop αSMA-positive SMCs, and overexpressing any of Notch1, 2, and 3 intracellular domains (ICD) can induce the SMC contractility genes in a serum response factor-dependent manner (SRF) (10, 38, 39). Thus, it is likely that the combined activity of Notch2 and Notch3 efficiently boosts the contractile differentiation. Following this notion, Notch2 and Notch3 silencing both downregulated pro-contractile genes OLFM2(40, 41), SPON2(42), and FOXS1, all of which were among the top 10 DEGs induced by Jagged1 at 24h (Fig. S1E). Especially, the Forkhead box family gene FOXS1 was highly responsive to Jagged1. It was regulated only by Notch2 at 4h, and both by Notch2 and Notch3 at 24h, similar to the putative Notch targets. FOXS1 is a periarterial marker(43, 44) and regulates TGFβ-mediated ACTA2 expression(45). Consistently, ChEA3(46) predicted FOXS1 as one of the top candidates upstream of *cluster d* (Fig. S1F). Thus, we tested TGFβ signaling and FOXS1 as potential factors favoring contractile differentiation downstream of Jagged1-induction. Silencing FOXS1 led to a non-significant but consistent decrease in ACTA2 and CNN1 expression. Meanwhile, SRF silencing abolished their expression altogether (Fig. S1G). Jagged1 induction or DAPT treatment did not affect SMAD2 phosphorylation. Additionally, inhibiting TGFβ receptor signaling did not affect Jagged1-induced αSMA production or collagen I or IV expression (Fig. S1H-J). These findings suggest that Jagged1-Notch2/3 signaling directly promotes procontractile genes that act upstream of SRF.

### Upregulation of Notch3 by the Jagged1-Notch2 axis contributes to prolonged Notch signaling

Considering the delayed contribution of Notch3 to the Jagged1-induced transcriptional response, we sought to understand the temporal relationship between Jagged1, Notch2, and Notch3 better. As Notch3 was already shown to be positively regulated by the Notch signaling(10, 47), we attributed its increased influence at 24h to its upregulation. Indeed, NOTCH3 mRNA was upregulated more than 2-fold upon ligand induction, whereas NOTCH2 mRNA seemed unchanged relative to the basal conditions (Fig. S1A). Interestingly, the Jagged1-induced expression of NOTCH3 was nearly lost upon Notch2 silencing. Similarly, the S1-cleaved Notch3 products were significantly downregulated with Notch2 silencing after 72 hours of Jagged1 induction (Fig. S2A,C), suggesting that the activation of the basal Notch3 was not sufficient to potentiate its own expression and that this process instead relied on Notch2 activity. To confirm this causal relationship, we performed DAPT washout experiments, where we grew SMCs on immobilized J1-Fc in the presence of DAPT overnight and removed the inhibitor the next day to allow Notch signaling activation. This way, we also hoped to eliminate any bias on the receptor cleavage that might arise from trypsinization at the time of cell plating on the ligands. First, we checked the protein abundance using the wild-type cells (Fig. 2A,B) and confirmed that the signal from the S1-cleaved Notch3 products increased by more than 5-fold starting from 24h. The corresponding bands for Notch2 were reduced by nearly 50% compared to the starting conditions, suggesting that Notch2 might be consumed at a faster rate, possibly due to the absence of similar positive feedforward mechanisms for this receptor. Next, we included SMCs with Notch2 or Notch3 silencing in a similar setting (Fig. 2C). The qPCR results recapitulated the same trends, showing that the effect of Notch3 on Notch target gene expression (HEY1 and JAG1) was negligible at 4h, but became more prominent at 24, consistent with the nearly 3-fold increase of the NOTCH3 mRNA following the Jagged1-induction. Again, the upregulation of NOTCH3 was nearly lost with Notch2 silencing, confirming our findings from the RNAseq.

**Figure 2.**
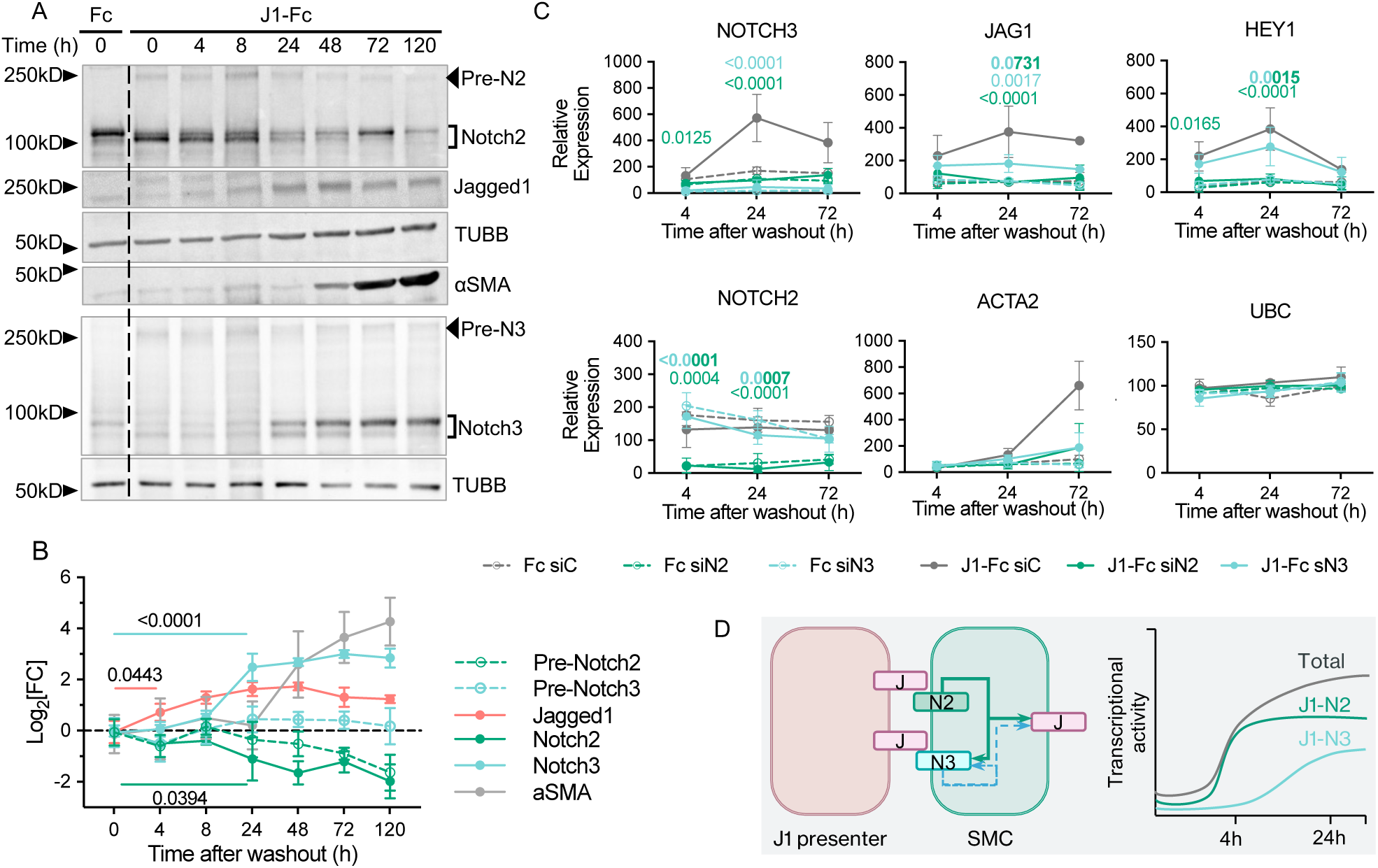
Upregulation of Notch3 by the Jagged1-Notch2 axis contributes to prolonged Notch signaling. (A, B)SMCs were grown overnight on immobilized J1-Fc in media containing 10μM DAPT. Following DAPT washout the next day, the cells were sampled at the indicated time points. The immunoblot analysis (A) and the quantified signal from the indicated proteins relative to the protein loading are shown (B). Pre-Notch is the full-length precursor before S1-cleavage. The dotted lines represent log2 fold change (mean±SD, n=3 independent experiments, one-way repeated measures ANOVA, and Dunnet’s test for multiple comparisons. Pairwise comparisons with p values smaller than 0.1 are indicated on the plot). (C) SMCs were transfected with the indicated siRNAs. 24h post-transfection, the cells were plated on immobilized J1-Fc and grown in media containing 10μM DAPT. Following DAPT washout the next day, the cells were sampled at the indicated time points. The dotted lines represent relative expression (mean±SD, n=3 independent experiments (n=2 for 72h). Two-way ANOVA and Tukeýs test for multiple comparisons were performed individually for 4- and 24-hour time points. Pairwise comparisons between J1-Fc-treated samples with p values smaller than 0.1 are indicated on the plots and color-coded for the groups compared. (D) Schematic description of the working model regarding the temporal relationship between Jagged1, Notch2, and Notch3 and the Notch signaling output over time.

To investigate the relevance of the upregulated Notch3, we also checked the abundance of Pdgfrβ, αSMA, the cell-intrinsic Jagged1, as well as the ACTA2, JAG1, and HEY1 mRNAs later at 72h. All molecules, except for Pdgfrβ, showed a negative trend upon Notch3 silencing (Fig. 2C; S2A,C). Contrary to the indications from the RNAseq, Notch3 did not mediate a visible reduction in SMC proliferation in the absence of Notch2 (Fig. S2B), suggesting that the initial levels of Notch3 might not be sufficient to potentiate this effect on its own. We next prepared spheroid cultures using only SMCs or SMCs and endothelial cells (ECs) to eliminate extrinsic J1-Fc ligands and instead enforce the native cell-cell Jagged1-Notch signaling. At 72h, both spheroids showed a similar reduction in the immunostaining of Jagged1 with Notch2 and Notch3 silencing (Fig. S2D and E), supporting that Notch3 contributes to the long-term Notch signaling response. These results together suggest that the initial Notch signaling “push” by the Jagged1-Notch2 axis is necessary for Notch3 expression, which then contributes to the long-term Notch signaling during *in vitro* SMC differentiation (See Fig. 2D for the schematic description of the proposed mechanism).

### Elevated levels of Notch3 are required for transcriptionally meaningful canonical Notch3 signaling due to the weak Notch3-CSL interaction

Next, we wondered why Notch3 relied on amplified expression to produce a sufficient transcriptional response to Jagged1. NOTCH3 is expressed nearly twice as much as NOTCH2 in SMCs, already at the basal levels, eliminating the possibility that the difference in activity was caused by the initial levels (Fig. S3A). Evidence suggests that the interactions between Jagged1 and Notch3 rely on additional molecules in SMCs (48–50). Hence, we asked whether receptor availability to J1-Fc could contribute to the weaker signaling from the Jagged1-Notch3 axis. To test this, we used a saturating concentration (10nM) of pre-labeled soluble J1-Fc to stain live SMCs (Fig. 3A). Confocal imaging following the fixation of the stained SMCs showed that silencing of Notch2 and Notch3 caused a visible difference in the fluorescent puncta (Fig. 3B).

**Figure 3.**
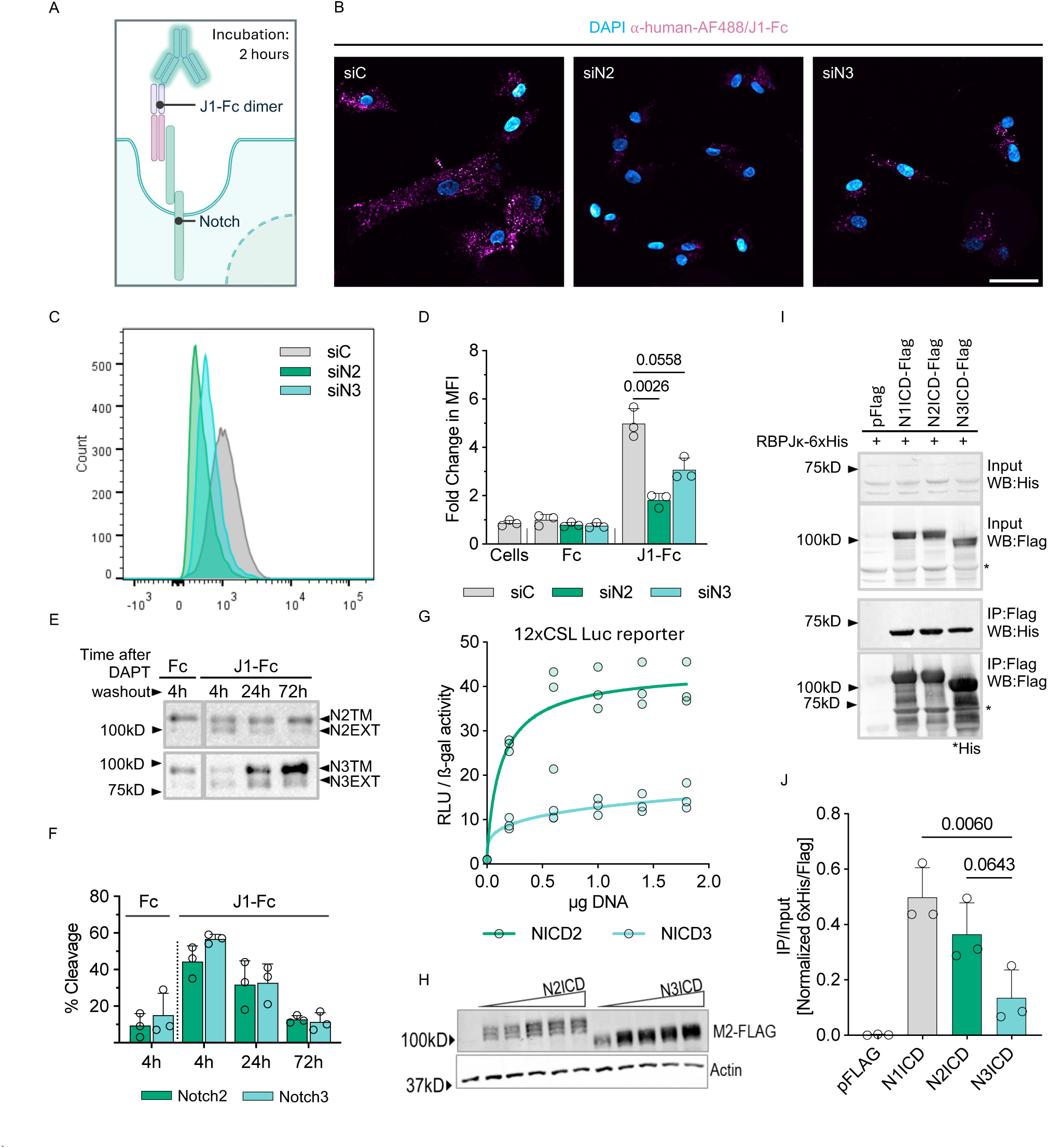
Elevated levels of Notch3 are required for transcriptionally meaningful canonical Notch3 signaling due to the weak Notch3-CSL interaction. *(A-D) SMCs were live-stained with fluorescently labeled J1-Fc 48h post-transfection with the indicated siRNAs to assess ligand-receptor interaction (A). Confocal slices (B) of formalin-fixed SMCs (control, Notch2KD, and Notch3KD) (Scale bar=50μm) and histograms (C) depicting the intensity histograms of the SMCs based on flow cytometry. The bar plots (D) show the fold change in mean intensity of the live cells stained with J1-Fc measured using flow cytometry (mean+SD, n=3 independent experiments, one-way ANOVA followed by Šídák’s test for multiple comparisons. Pairwise comparisons between J1-Fc-stained samples with p values lower than 0.1 are shown on the plot.)*. *(E, F) Immunoblots produced by C-terminal probing of Notch2 and Notch3 following DAPT washout in SMCs plated on immobilized J1-Fc, and the quantitative analysis of the bands. The receptor cleavage was measured by the ratio of the NEXT (lower band) to the total S1-processed fragment (NEXT+TMIC (higher band)) (mean+SD, n=3 independent experiments, one-way ANOVA followed by Šídák’s test for multiple comparisons. Pairwise comparisons with p values lower than 0.1 are shown on the plot.)*. *(G,H) CSL-dependent Notch transcriptional activity in HEK 293T cells transfected with increasing amounts of Flag-tagged Notch2 or Notch3 ICD and 12xCSL-luciferase (G). The luciferase activity is presented as luminescence intensity relative to β-galactosidase activity (mean+SD, n=3 independent experiments. Data are presented as regression curves and circles as replicates.). (H) Immnunoblots indicating efficient transfection of the NICD-flag constructs*. *(I, J) Flag-tagged NICD paralogs were immunoprecipitated from HEK 293T cells co-expressing 6xHis-tagged RBPJκ (I). The Flag and the 6xHis IP signals were normalized to the input. The normalized 6xHis/Flag signal is presented in the bar plot (J) (mean+SD, n=3 independent experiments, one-way ANOVA followed by Tukey’s test for multiple comparisons. Pairwise comparisons between NICD paralogs with p values lower than 0.1 are shown on the plot.)*.

We repeated the live staining protocol and measured the fluorescence signal using flow cytometry. Again, silencing either receptor caused a comparable decrease in the fluorescence signal (Fig. 3C,D). We observed a marginally larger, albeit statistically insignificant, decrease in the mean fluorescence intensity of the Notch2-silenced cells. We attributed this difference to the slight reduction of Notch3 observed with the Notch2 siRNA. Thus, we concluded that the interaction of J1-Fc with Notch2 and Notch3 was comparable. We also investigated receptor processing triggered by the immobilized J1-Fc at 4, 24, and 72 hours post-DAPT washout due to the evidence that the Notch paralogs vary in sensitivity to cleavage (51–54). We did not find meaningful differences between Notch2 and Notch3 processing, measured by the ratio of the NEXT to the total signal from the NEXT and the NTM bands (Fig. 3E,F). These results together suggested that the large difference in the transcriptional activity of the two receptors could not be explained by J1-Fc binding preference or the differential regulation of receptor processing. We next asked whether the intracellular domains of the two receptors elicited different transcriptional activity. We tested this by expressing increased concentrations of NICD2 and NICD3 constructs bearing Flag tags, together with a 12xCSL-Luciferase reporter and β-galactosidase in HEK 293T cells (Fig. 3G,H). NICD2 mediated dramatically higher reporter activity, reaching far beyond the dynamic range of NICD3 even at very low concentrations. To test the interaction between the NICD paralogs and RBPJκ, we co-expressed 6xHis-tagged RBPJκ and each of the three NICD paralogs (NICD 1, 2, and 3) in HEK 293T cells and performed co-immunoprecipitation using beads bearing α-Flag antibody. Western blot and densitometry analyses showed that NICD1 and NICD2 interacted more strongly than NICD3 with RBPJκ (Fig. 3I,J).

Next, we used computational modeling to ask if the difference in transcriptional activity mediated by Jagged1-Notch2 and Jagged1-Notch3 signaling in SMCs can be explained with the NICDs transcriptional activity. The model, extended from previous versions (13, 14) to include both Notch2 and Notch3 signaling (Fig. S3B,C), could successfully replicate the qPCR data from Fig. 2G (the data at 24h), but only when assuming that NICD2 had a much higher potential to drive JAG1 and NOTCH3 expression than NICD3. Based on the optimization, the algorithm predicted that NICD2 would exert a significantly larger influence, more than 2-fold greater, on the production rates of Notch3 and Jagged1 compared to NICD3 (Fig. S3D-F). Our lab has previously shown that the RAM domain of NICD1 has a much lower dissociation constant (K_d_=0.022 μM) than NICD3 (K_d_=0.187 μM) for RBPJκ, implying higher affinity between NICD1 and RBPJκ(55). NICD1 and NICD2 are structurally similar and have been shown to be functionally interchangeable during murine development and carcinogenesis(51, 56). Thus, these results suggest that the weak transcriptional consequence of the Jagged1-Notch3 axis is caused by the low potency of the Notch3 ICD to drive RBPJκ-mediated transcription, and that higher levels of Notch3 are needed for meaningful Notch signaling output during SMC differentiation.

#### Levels of Notch3 predict artery-typical vulnerability to altered Jagged1-Notch2/Notch3 signaling in the SMCs

Vascular SMCs show remarkable heterogeneity in their transcriptional profile, which contributes to region-specific vulnerabilities to perivascular disease (57–59). Likewise, Notch3 expression in the outflow tract has been suggested to depend on the embryonic lineage, where the SMCs of the neural crest origin show higher Notch3 expression than those derived from the sclerotome and the second heart field(25). Therefore, we wondered whether possible regional differences in the expression of Notch components and their differential ability to induce canonical Notch signaling could help explain receptor-specific effects in perivascular disease onset. We started by mining the expression of Notch signaling components in the arterial cells from multiple locations using previously published single-cell RNAseq data from adult mice (35, 57, 60) (Fig. 4A; S3I,H). Notch2 was expressed at relatively low levels in all arteries and showed little variation. Surprisingly, Notch3 expression varied vastly among the SMCs of different arteries. It was expressed highly in smaller muscular arteries, such as the arteries in the heart, the brain, and the colon. Meanwhile, the aorta, the proximal and the distal regions combined, had strikingly little Notch3 in comparison.

**Figure 4.**
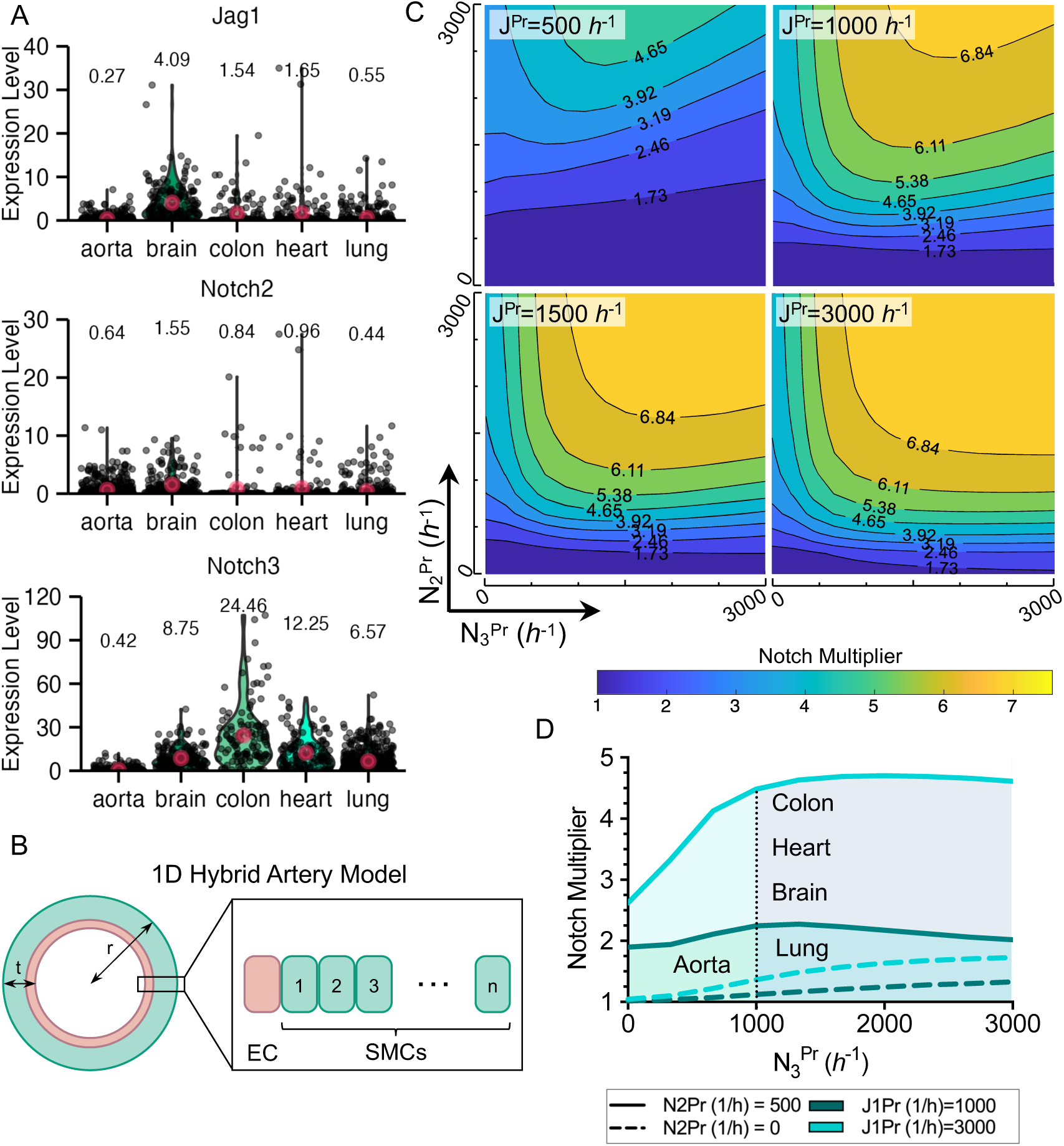
Levels of Notch3 predict artery-typical vulnerability to altered Jagged1-Notch2/Notch3 signaling in the SMCs. (A) Normalized expression of Jag1, Notch2, and Notch3 in arterial SMCs of different mouse organs extracted from scRNAseq data (Muhl et al.(35); aorta, heart, colon, and lung) and (Vanlandewijck et al.(57); brain). The circles and the text above the violin plots denote the mean expression. (B) Schematic one-dimensional array of 1D artery model with an endothelial cell and a row of SMCs. The cascade of signaling is initiated by constant Jagged1 emission from the endothelial cell (EC) and SMCs are interacting with their neighboring cells. (C) Heatmaps showing the simulated impact of the initial Jagged1, Notch2, and Notch3 levels on the Notch signaling readout (Notch multiplier) in the 1D hybrid arterial wall. (D) 2D projection of the predicted ranges of Notch multiplier in arterial SMCs of different organs with respect to their stoichiometrically scaled levels of Jagged1, Notch2, and Notch3 based on A, and in a Notch2 knockout scenario.

Next, we aimed to see how expression of the two receptors could affect the Notch signaling output in the arterial wall. Using the optimized parameters relating to the Jagged1-Notch2/3 signaling from our simulations (Fig. S3F - Also see Table S2 in Supporting Methods), we extended the computational model of Notch signaling in the arteries that was previously adapted and tested by our lab (13, 61–63) (Fig. 4B). The new model takes Notch2 and Notch3 dynamics during Jagged1- mediated lateral induction into account and calculates a “Notch multiplier” (Notch multiplier=1 indicates basal Notch activity) as a measure of the Notch signaling activity in a simulated arterial wall. We then adjusted the basal Jagged1, Notch2, and Notch3 production rates in SMCs 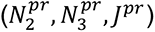 in a range between 0 and 3000 ℎ^−1^ to predict the Notch multiplier in different Notch2/Notch3 stoichiometries (Fig. 4E). The model was highly sensitive to Jagged1 and Notch2 manipulations, as the Notch multiplier rapidly changed with respect to 𝐽^𝑝𝑟^ and 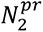. This is also consistent with the previously reported *in vivo* findings, where combined heterozygous Jag1^ΔDSL^ and Notch2^del1^ mutations in mice closely recapitulate the ALGS phenotype (64). Meanwhile, increasing 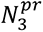 by itself had little impact on the Notch multiplier and its contribution was merely additive to that of 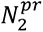, corroborating our *in vitro* data.

To put the Notch multiplier in a physiological perspective, we chose Notch stoichiometries resembling those in different arterial SMCs. Considering the low expression of Notch2 and the relatively higher Notch3 expression in all arterial SMCs, we assigned simple Notch2^Low^/Notch3^Low^ 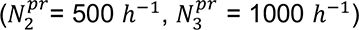 and Notch2^Low^/Notch3^High^ stoichiometries 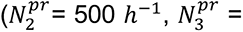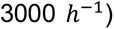 for aortic and pulmonary SMCs, respectively. We set the minimum 𝐽^𝑝𝑟^ to 1000 ℎ^−1^, since otherwise, the Notch multiplier was not meaningful with the chosen ranges of 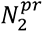 and 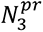. For the muscular arterial SMCs (heart, colon, and brain arterial SMCs), we used the same Notch2^Low^/Notch3^High^ stoichiometry but set 𝐽^𝑝𝑟^ to 3000 ℎ^−1^ in line with the higher Jag1 expression seen in these cells. This way, we could fit the numbers representative of different arterial SMCs within the dynamic range of our simulations (Fig. 4D). This approach yielded Notch multipliers resembling the trend of Nrarp expression, a proxy for Notch activity, across different arterial SMCs (Fig. S3H). Furthermore, setting 𝐽^𝑝𝑟^ to 0 with the aortic receptor parameters reduced the Notch multiplier from 2.24 to nearly 1 (Fig. S3G), resulting in a similar negative 2-fold change that was seen with Notch target expression in the mouse aortic smooth muscle tissue upon Jagged1 knockout (65).

Further, we explored loss-of-function scenarios for Notch2 or Notch3 by setting the production rates of either to 0. The model predicted that Notch2 knockout, but not Notch3 knockout, reduces the Notch multiplier in all arterial conditions significantly (Fig. 4D). Setting 𝑁^𝑝𝑟^ to 0 resulted in an approximately 2-fold decrease in the Notch multiplier in the aortic SMCs (2.24 to 1.12), leading to almost a complete loss of the Notch activity. Although the pulmonary SMCs expressed three times as much Notch3, the decrease in the Notch multiplier with Notch2 knockout was comparable with the aortic SMCs (2.02 to 1.33), consistent with the importance of Notch2 in the outflow tract. The Notch multiplier in the muscular arterial SMCs, which express three times as much Notch3 and Jag1 as their aortic counterparts, was reduced from 4.61 to 1.73 with Notch2 knockout. However, we observe that 1.73 is reasonably close to the Notch multiplier that the model predicted for the aorta and the pulmonary arteries (approximately 2) in the basal conditions. Therefore, one can argue that this level of Notch activity might be sufficient during SMC development. These results suggest that elevated expression of Notch3 can induce sufficient canonical activity, provided high basal Jagged1 expression in the SMCs. Since periarterial lateral induction is triggered by the EC-bound ligands (12, 66), we checked the ligand expression in ECs from the same organ datasets. The ECs of the muscular arteries did not show elevated Jag1, unlike the SMCs (Fig. S3I). However, ECs express high levels of Dll4, which has been shown to contribute to the signaling between the coronary endothelium and the developing SMCs (67). Thus, Dll4 and Jagged1 together might induce sufficient Notch signal in the first layer of SMCs to kickstart lateral induction in the developing media. Supporting this notion, mice with SMC-specific knockout of Jagged1 develop an SMC layer adjacent to the endothelium in the descending aorta, but this layer is much thinner compared to the wildtype, highlighting the importance of Jagged1 of the SMC-origin (68).

Overall, our model predicts that Notch3 depletion has a negligible impact on the periarterial Notch signaling in the presence of Notch2, and that Notch2 depletion is highly detrimental, especially for the aortic-pulmonary tree. On the other hand, it predicts that expression of Notch3, particularly in smaller muscular arteries where it is expressed highly, can partially compensate for Notch2 depletion, but the compensatory effect of Notch3 is dependent on the complementary expression of Jagged1, highlighting the importance of this ligand and its expression level during periarterial development.

### Developing SMCs at the cerebral base can compensate Notch2 depletion in the zebrafish larvae

We tested if we could capture the artery-typical outcomes for impaired Notch2 and Notch3 signaling predicted by our model, in an *in vivo* developmental setting. For this purpose, we chose zebrafish larvae, an established model for vascular development (69). Zebrafish larvae with Notch2 and Notch3 deletion were shown to have a reduced number of pdgfrb^High^ early mural cells in the dorsal aorta (DA), and the cerebral base arteries, BCA (basal communicating artery) and PCS (posterior communicating segment), respectively (9). Thus, we started by assessing the Notch receptor expression in the developing zebrafish perivascular cells in these locations (Fig. 5A). There are no reliable antibodies for the zebrafish Notch orthologs; thus, we performed *in situ* Hybridization Chain Reaction (HCR) in 3dpf (3 days post-fertilization) larvae expressing a pdgfrb reporter using probes against notch2 and notch3. We could not reliably quantify the total intensity or the number of mRNA puncta per cell, due to the difficulty of segmenting the pdgfrb^High^ cells and the irregular signal from notch3. However, notch3 stained more consistently and was localized in broader pdgfrb+ areas in the cerebral base, while the staining was sparser in the dorsal aorta. On The signal from notch2 looked similar in both regions and appeared as sparse puncta (Fig. 5B,C).

**Figure 5:**
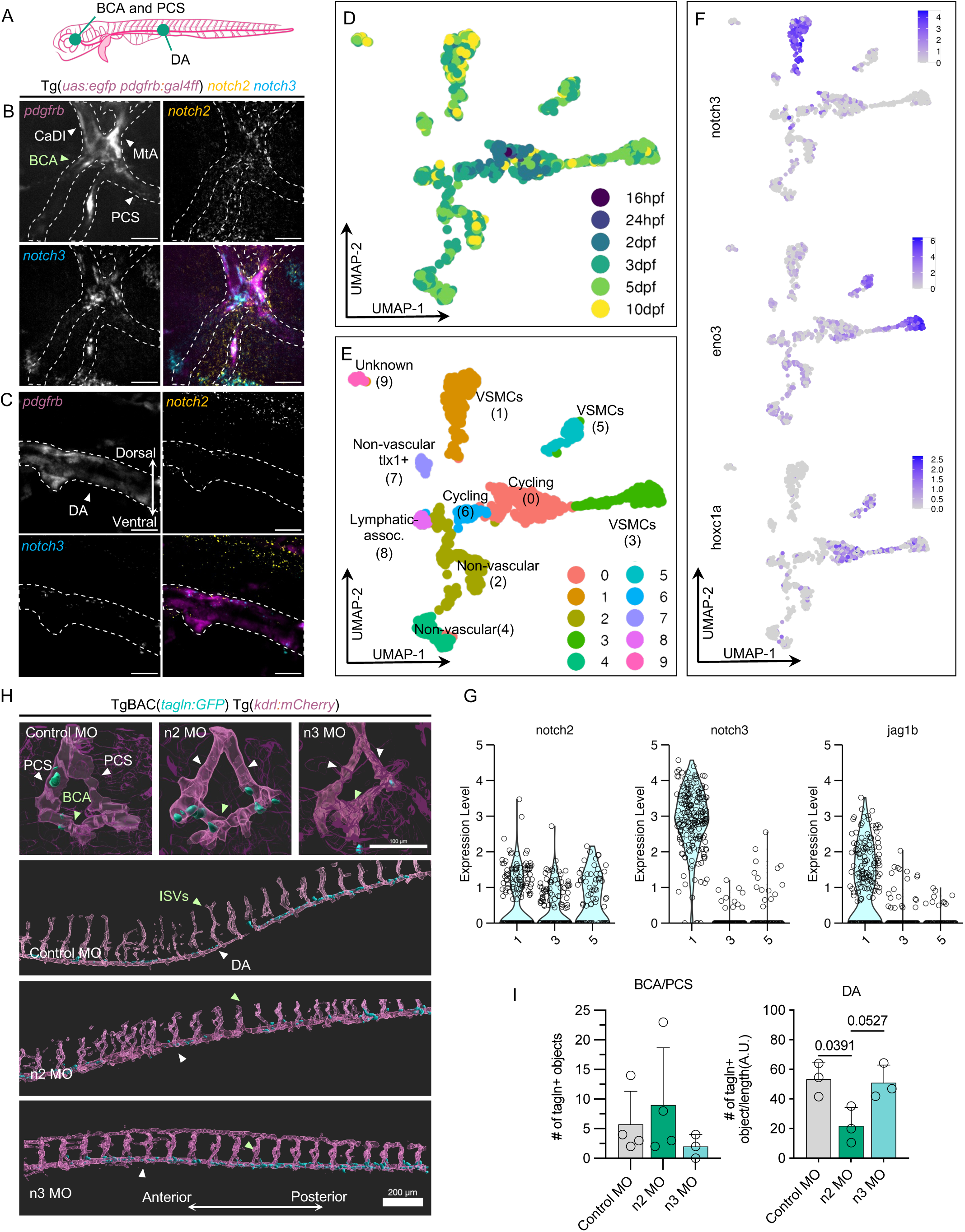
Developing SMCs at the cerebral base can compensate Notch2 depletion in the zebrafish larvae. (A) Schematic description of the vascular tree of the zebrafish larvae, and the position of the DA, the PCS, and the BCA (Created using Biorender). (B, C) In situ RNA HCR was performed with probes against notch2 and notch3 in 3dpf zebrafish larvae bearing a pdgfrb reporter. Confocal sections of the arteries from the cerebral base (B) and the dorsal aorta (C) given with dashed lines loosely marking the arterial regions based on the pdgfrb signal are shown (Scale bars=20 μm) (DA: dorsal aorta, BCA: basal communicating artery, PCS: posterior communicating segment, CaDI: caudal division of the internal carotid artery, MtA: metencephalic artery) (D, E) UMAP projection of the SMCs color-coded based on the Seurat clusters and the developmental stage from the Zebrahub mesoderm scRNAseq dataset. Cluster annotations were based on known markers of vascular and non-vascular SMCs as well as notch3 expression (Please also see Fig. 4S for the markers used for annotation). (F, G) Normalized expression of notch3, trunk marker eno3, and anteroposterior position-related homeobox gene hoxc1a (Color gradient denotes log-normalized expression.); and (G) the normalized expression of the Notch signaling components in the indicated vascular SMC clusters. (H, I) The embryos bearing a contractility reporter (tagln:GFP) or the endothelial reporter (kdrl:mCherry) were injected with 10ng control, notch2 or notch3 morpholinos. The orthogonal view of the surface-rendered light sheet images of the indicated arteries from the 102 hpf (H, the BCA/PCS, scale bar=100μm and H, the DA and the ISV, scale bar=200μm). The number of tagln+ objects associated with the endothelium are shown in the bar plots (I) (mean+SD, n=3 or 4 embryos per morphant group, one-way ANOVA followed by Tukeýs test for multiple comparisons. Pairwise comparisons with p values smaller than 0.1 are shown on the plots.).

For a more quantitative approach, we turned to a single-cell RNAseq dataset from Zebrahub (70). Here, we chose to look at the mesoderm dataset, since mural progenitors at the cerebral base and the dorsal aorta mainly derive from this lineage (71). We extracted the cell cluster, compiling all SMCs under the annotation “visceral muscle”. We then created UMAP projections using the Seurat package (72) (Fig. 5D) and identified the vascular SMC clusters by sorting out the cells expressing non-vascular SMC markers desmb, nkx2.3, rem1, and tlx1 (Fig. S4A). Among the remaining, the clusters #1, #3 and #5 were positive for the SMC markers acta2, tagln, and myh11a (Fig. S4). Out of these clusters, cluster #1 expressed dramatically higher levels of notch3 compared to the others (Figure 5F,G). These cells are enriched for the head SMC marker foxc1 (73) and pericyte-associated signatures (pdgfrb, zeb2a, rasl12, and cd248a) (Fig. S4A) consistent with the more pericyte-like properties of the SMCs in the smaller cerebral (74) and coronary arteries (26), and cluster closely with pharyngeal arch and periocular mesenchyme cells in the original dataset (Fig. S4B,C). Moreover, unlike the other SMC clusters, these cells were largely negative for spatially restricted homeobox genes (e.g., hoxc1a and hoxc8a), somite/trunk signatures (eno3 and fbp2), and fibroblast-like markers (pdgfra and tgfbi) (Fig. 5E; S4A) (75), suggesting that they are from vessels anterior to hindbrain. They also showed higher levels of jag1b and a notch2^Low^/notch3^High^ stoichiometry (Fig. 5G), recapitulating the same expression pattern as the murine brain SMCs.

Finally, to test whether elevated notch3 could drive the differentiation of these cells without the need for notch2, we injected embryos bearing tagln and kdrl reporters with notch2 or notch3 morpholinos. Light sheet microscopy at 102 hpf (hours post-fertilization) revealed that the notch2 morphants had a significantly decreased number of tagln+ objects around their dorsal aorta, whereas the notch3 morphants were indistinguishable from the control fish, supporting the negligible contribution of notch3 to SMC differentiation in this location (Fig. 5H,I). At BCA and PCS, the number of could not detect any differences between the notch2 morphants and the control. However, consistent with the loss of the pdgfrb^High^ cells in this location with notch3 knockout (9), the notch3 morphants showed a negative trend in the number of tagln+ objects (Fig. 5H,I). Overall, these results suggest that developing zebrafish SMCs with elevated Notch3 expression could tolerate Notch2 depletion during SMC differentiation, corroborating our simulations.

## DISCUSSION

Notch signaling is integral in perivascular cell specification, maturation, and homeostasis. As a result, its dysregulation is linked to congenital and acquired diseases with catastrophic consequences for the vasculature. Here we studied the signaling mechanism governing Jagged1 and the receptors Notch2 and Notch3, and how it regulates arterial SMC development. Our findings support a causal role of Notch2 for Notch3 (10). We show that when induced with Jagged1, Notch2 and Notch3 both regulate genes related to vascular SMC specification, maturation, and quiescence, albeit with lower transcriptional activity by Notch3. This supports the previous notion that the two receptors could essentially be mutually compensatory during SMC development (9, 10). We cannot state that Notch2 and Notch3 specifically regulate different genes without NICD-DNA interaction assays, such as ChIPSeq, especially given the temporal relationship between Notch2 and Notch3. Thus, we do not exclude the possibility that they might otherwise have specific targets that are not directly related to SMC differentiation, considering the genes associated with the pulmonary-aortic tree regulated by Notch2 in our RNAseq data. As for Notch3, we could not detect any specific signatures among the Jagged1-induced genes that would suggest a pro-growth/survival function as previously suggested (30, 76). Therefore, the canonical Jagged1-Notch3 signaling does not directly explain pathological SMC growth related to increased Notch3 activity or decreased SMC survival upon Notch3 loss. Instead, Notch3 might have paradoxical functions due to non-canonical mechanisms independent of Jagged1. Indeed, it was shown to go through ligand-independent cleavage to promote cell growth in basal breast cancer (77) and signal non-canonically by interacting with β-catenin to promote cell survival in lung cancer (78). Considering that Notch3 itself is upregulated as a result of Jagged1-Notch signaling, in a perivascular context, one must carefully consider whether a particular effect is linked directly to the Jagged1-Notch3 axis or the enhanced cell-autonomous functions of Notch3 due to its increased abundance.

We show that the Jagged1-Notch3 axis is weaker in inducing a transcriptional response compared to Notch2 and relies on the high abundance of both Jagged1 and Notch3. This has the following implications for the Notch-related vascular disease context. First, the low canonical activity of Notch3 would imply a heavier burden on the Jagged1-Notch2 axis during smooth muscle cell development than previously thought, especially for those in the aorta, where Notch3 is expressed in lower quantities compared to the smaller arteries. This is consistent with the severe deformations observed in the outflow tract of mice with Notch2 and/or Jagged1 depletion, such as patent ductus arteriosus (24) and aortic narrowing (23). Similarly, ALGS patients frequently present with narrowing defects in the outflow tract, such as pulmonary stenosis and aortic coarctation (16, 18). It is worth mentioning that, unlike a severe Notch2 depletion scenario we explored in our experiments, Notch2 mutations reported in the ALGS patients are often heterozygous missense mutations that would lead to a moderate reduction in the receptor activity. Consistently, while total Notch2 knockout alone is embryonically lethal in mice on E10.5(79), the hypomorphic Notch2^del1/del1^ mutation can be partially rescued by Notch3 even in the developing mouse aorta (10). This might help explain why Notch2-type ALGS is not as strongly associated with vascular defects as are the Jagged1-type cases. Other evidence suggests that the induced knockout of Notch2 in adult mice does not affect the neointimal area following carotid injury (80) or diet-induced atherosclerotic plaque formation (81). However, Notch2 activity might be more integral during the development of SMCs than in their homeostasis or injury response, since in the latter case, the perivascular tissue is already established and rich in supporting extracellular matrix that can switch on compensatory mechanisms. Indeed, in the absence of elastin, epigenetic mechanisms boost Jagged1-Notch3 signaling in the aorta (31). Similarly, carotid injury leads to transient expression of Notch1, which contributes to injury response (82).

Second, due to the low transcriptional contribution of Notch3, our results indicate that perivascular defects related to the dysregulation of the Jagged1-Notch3 axis (e.g., in CADASIL) might be associated with the sustainment of the differentiated SMC state and homeostasis, rather than early SMC development. Consistently, the late contractile SMC marker Myh11, but not the SMC precursor emergence, is affected by Notch3 depletion in the mouse heart (26). Additionally, there is growing consensus that the molecular mechanism underlying the poor SMC coverage in CADASIL is the abnormal accumulation of the Notch3 ECD on the cell surface due to the obstruction of its ligand-dependent elimination, pointing at a gain-of-function effect of Notch3 rather than defective canonical Notch3 signaling (83–86). A Jagged1 mutation in mice phenocopying the human ALGS (Jag1^Ndr^) shows partial loss in its binding capacity to Notch3 ECD(87) and causes poor SMC coverage in the retinal arterioles and the coronary arteries with increased SMC apoptosis (88) similar to CADASIL, asking whether this phenotype links to a defect in the elimination of the Notch3 ECD. Here we show that smaller muscular arteries like those in the brain and the heart express elevated levels of Notch3, which could boost Notch3 ECD accumulation in these locations and contribute to the small vessel specificity of CADASIL. Indeed, mutant Notch3 (R169C) does not cause a change in the expression of Notch3-driven genes in the arteries, but the forced expression of two copies of the mutant in addition to the wild-type Notch3 exacerbates the ECD accumulation and the resulting arteriopathy. Meanwhile, eliminating one copy of the wildtype Notch3 alleviates the phenotype (89), supporting a Notch3 overdose mechanism over a Notch3 hypo-signaling scenario. Additionally, we have previously shown that Notch3 and Jagged1, but not Notch2, are negatively regulated by strain (13, 90). Thus, it is likely that altered hemodynamics, like high blood pressure, has a considerable impact on how Notch3 functions in smaller arteries during disease. Overall, the differences in the canonical activity of Notch2 and Notch3, and the artery-typical variations in their stoichiometry, can partly explain the locally restricted periarterial phenotypes and clinical findings relating to dysregulation of Jagged1, Notch2, and Notch3.

## LIMITATIONS OF THE STUDY

This study has several limitations. Most importantly, our findings are based on the mRNA level expression, and thus, we cannot dismiss the possibility that Notch2 and Notch3 are regulated differently at the protein level. Indeed, contrary to our predictions, Notch3 knockout alone causes an almost complete loss of perivascular cells in the cerebral base of zebrafish larvae (9), showing that Notch2 cannot compensate for Notch3 depletion in this location. While we found that all vascular SMC clusters in zebrafish larvae expressed comparable levels of Notch2 based on Zebrahub, in reality, only a small fraction of the cells in the cerebral arteries were found to be positive for Notch2 reporter activity(9). Thus, we acknowledge the need for improved tools to assess different Notch protein levels *in vivo* and say whether some cells might rely solely on Notch3. Additionally, Notch targets respond differently to the temporal abundance of NICD (91) and might be regulated by co-transcriptional factors in addition to RBPJκ (92). Indeed, cerebral-enriched Foxc1 boosts NICD activity in the zebrafish (93). Notch3 might activate contractility-related genes more efficiently than the other putative Notch targets, as implied by our RNAseq data. Thus, we cannot exclude the possibility that the transcriptional influence of Notch3 might be higher than our predictions. Finally, we did not include the possible influence of Notch1 in our model, which was lowly expressed in our cultured cells as well as the scRNAseq datasets we investigated. Notch1 knockdown was shown to not affect periarterial cell emergence in the zebrafish trunk or brain(9). Similarly, heterogeneous Notch1 knockout does not affect SMC markers(94). However, we cannot rule out a possible compensatory effect from Notch1 in the absence of Notch2 or Notch3.

## AUTHOR CONTRIBUTIONS

SS, MS, and CS conceptualized the work. SS, NV, HH, and TR designed the experiments. SS, NV, HH, and TR executed the experiments, analyzed the data, and prepared the figures. SS wrote the initial draft. SS, NV, HH, TR, MS, SL, and CS revised and edited the draft. TR, MS, SL, and CS supervised the study. TR, SL, and CS acquired funding for the study.

## ACKNOWLEDGEMENTS

The bulk RNA sequencing and the data analysis were done in the Finnish Functional Genomics Center and the Medical Bioinformatics Center, Turku. All zebrafish work was done in Zebrafish Core Facility in Turku Bioscience. All microscopy experiments were done in the Cellular Imaging Core in Turku Bioscience. We would like to thank previous laboratory trainees Amanda Norren and Ana Ayala Perez for their help with the differentiation assays. We also thank other persons for the following: help with laboratory and administrative work (Alexandra Manea), help with the design of the RBPJκ plasmid (Elenaé Vázquez-Ulloa), guidance and help with flow cytometry assays (Ezgi Özliseli), guidance and help with qPCR assays (Marjaana Parikainen and Kati Kemppainen), guidance and help with light sheet microscopy and image analysis (Markus Peurla and Anna-Mari Haapanen-Saaristo), guidance with zebrafish experiments (Ilkka Paatero), and help with setting up the computational model (Margot Passier).

## Methods

Cell culture, Western blotting, RNA extraction, cDNA synthesis and qPCR analysis, microscopy specimen preparation, flow cytometry, re-analysis of the published single-cell RNAseq data, as well as Next Generation Sequencing of bulk transcriptome and data processing, and computational modelling of extrinsic ligand-induced Notch signaling are described in Supporting Methods. All reagents and resources used in this study are detailed in Table S4.

### Notch trans-activation assays

The cells were plated on the immobilized ligands in SmGM^®^-2 (Lonza Cat. #CC-3182). In all experiments, human IgG_1_ Fc (Bio-techne Cat. #110-HG) was used as a negative control. For the DAPT washout experiments, the cells were plated on the ligands in media containing 10μM DAPT ((2S)-N-[(3,5-Difluoro phenyl)acetyl]-L-alanyl- 2-phenyl]glycine 1,1-dimethyl ethyl ester) (Bio-Techne Cat. #2634-10) or DMSO (dimethyl sulfoxide) (Merck Cat. #D2650) as the vector control where appropriate and let recover for 24 hours. The next day, the DAPT was removed by replacing the media with fresh, warm media, and the cells were incubated for the indicated time until use. For the TGFβR inhibition experiments, the cells were plated on the ligands with media containing 10μM SB-431542 (MedChemExpress, Cat. #HY-10431), 10μM DAPT, or DMSO and incubated for either 24 hours for qPCR or 72 hours for immunoblotting. The cells were supplied with media containing fresh dilutions of the inhibitors every 24 hours and washed once with cold HEPES-buffered saline (HBS) (Lonza Cat. #CC50-34) before lysis.

### Ligand binding assays

The live Notch staining assay was done as described previously(95) with a few adjustments. 10^4^ SMCs per cm^2^ transfected with siRNAs against Notch2 and Notch3 were plated 24 hours post-transfection and left to incubate for another 24 hours. The next day, the cells were blocked with sterile-filtered SmGM^®^-2 containing 1% BSA and 10% goat serum (Biowest Cat. #S2000) for 45 min at 37°C with 5% CO_2_. 50nM Jagged1-Fc was conjugated with anti-human Alexa fluor 488 antibodies (Thermo Fisher Scientific Cat. #21091) at a 1:200 dilution in PBS for 1 hour at cold. The ligand dilution was added directly to the blocking media at a 1:5 ratio to achieve a final ligand concentration of 10nM and incubated for 2 hours at 37°C with 5% CO_2_. The cells were either used for microscopy following fixation or live for flow cytometry.

### Notch reporter assays

The luciferase assay was done as described previously(55). HEK 293T cells were transfected with 0.1μg 12xCSL-Luciferase (RBPJκ) and 0.1μg β-galactosidase along with increasing concentrations (0-1.8μg) of Flag-tagged Notch2 or Notch3 intracellular domain (NICD2 and NICD3) constructs. Each transfection was completed with 2μg using the plasmid, only bearing Flag. An extra set of cells was prepared in parallel to assess NICD abundance by immunoblotting. The next day, the cells were lysed in passive lysis buffer (Promega Cat. #E1941). The luciferase activity was measured using VivoGlo™ luciferin (Promega Cat. #P1041) and normalized to β-galactosidase activity.

### Bulk RNAseq

Total mRNA was isolated from SMCs treated with J1-Fc as described in Fig. 1A. The bulk RNA sequencing (RNAseq) was performed in the Finnish Cultural Genomics Center, University of Turku. Processing of the sequencing data analysis was done in the Medical Bioinformatics Center, University of Turku (See detailed pipeline in the SI Methods). Differentially expressed genes (DEGs) upon J1-Fc treatment in the control siRNA-treated samples were used for further analysis for each time point separately. For biological interpretation, z-scores for the DEGs were calculated based on 𝑙𝑜𝑔_2_(1+CPM) across all groups from each time point separately. The genes were then clustered using Ward’s method and used to build a heatmap using the pheatmap package (v.1.0.12). Gene ontology analysis for biological processes on each cluster was performed with the clusterprofiler package(v.4.14.4).

### Immunoprecipitation

HEK 293T cells expressing Flag-NICD and 6xHis-RBPJκ were lysed in RIPA buffer (50 mM Tris-HCl pH = 8.0, 1% Triton X-100, 1% sodium deoxycholate acid, 0.1% SDS, 1 mM Na4VO3, 50 mM NaCl, 10 mM NaF, and 1 mM EDTA) containing 1x cOmplete™ protease inhibitors (Roche, Cat. #05892970001) on ice for 10 min. The lysates were scraped into 1.5 ml Eppendorf tubes and centrifuged for 15 min at 14,000 RCF, 4°C. A total of 1.5 mg protein was immunoprecipitated with 40μl of anti-Flag affinity agarose beads (Merck, Cat. #A2220) on rotation for 1 hour at 4°C. The beads were washed with the lysis buffer three times and boiled in 3x Laemmli buffer for 10 minutes at 95°C.

### Zebrafish lines and reporter assays

All zebrafish work was carried out in the Zebrafish Core Facility at Turku Bioscience Center under license ESAVI/31414/2020 and ESAVI/44584/2023 granted by the Regional Administration Office of Southern Finland. The offspring of transgenic zebrafish heterozygous for tagln (tagln:GFP) or kdrl (kdrl:mcherry) were injected with 10ng control, Notch2, or Notch3 morpholinos (Gene Tools Store) before the 8-cell stage. Embryos were kept in E3 media (5 mM NaCl, 0.17 mM KCl, 0.33 mM CaCl_2_, 0.33 mM MgSO_4_) supplemented with 30 mg/ml of 1-phenyl 2-thiourea (PTU) (Merck, Cat. #P7629) at 28.5°C. At 4 dpf, the larvae positive for both reporters were mounted dorsally in 2% agarose molds, immobilized with 1% agarose, and imaged in the same media containing 160 mg/ml tricaine (Merck, Cat. #E10521) using MSquared Aurora light sheet microscope with 0.4 μm slice thickness. The stacks were stitched using the Zeiss Arivis Vision 4D software (v.3.5.) and analyzed using the Oxford Instruments Imaris image analysis software (v.10.2.0). Briefly, the endothelium marked with kdrl and the tagln+ objects were surface-rendered following pseudo-flat field correction for uneven illumination. The tagln+ objects that were at least in 5μm proximity of the endothelium were considered vascular-associated. In the hind/midbrain region, the objects that were deemed artifacts of the tagln+ tip of the notochord were manually chosen and left out of the analysis. The number of tagln+ objects in the dorsal aorta was normalized to the length of the measured region and presented in arbitrary units.

### *In situ* hybridization chain reaction (HCR)

3 dpf transgenic larvae expressing a pdgfrb reporter (UAS:EGFP, pdgfrb:GAL4FF) were euthanized in E3 media with tricaine at cold for 1h and fixed in 4% PFA overnight at cold. After five washes with PBS, the embryos were dehydrated and permeabilized in methanol at RT four times for 10 min and once for 50 min and rehydrated with four gradually decreasing concentrations of MeOH in PBS with 0.1% Tween 20 (PBST) for 5 min each. The larvae were treated with RNAScope™ Protease Plus (Biotechne, Cat. #322331) for 30 min at RT and quickly washed twice with PBS. The larvae were post-fixed in 4% PFA at RT for 20 min, washed five times with PBS for 5 min each, and used for HCR™ Gold RNA (Molecular Instruments) *in situ* hybridization reaction according to the suppliers’ protocol, adapted from Choi et al.(96). The notch2 and the notch3 probes were visualized using x2-546 and x1-647 hairpin pairs, respectively. The fish were mounted ventrally directly on 35 mm glass-bottom confocal dishes (VWR, Cat. #75856) in 1% agarose (w/v) and imaged by taking z-stacks using the 3i CSU-W1 spinning disc confocal microscope with the 25x/0.8 Imm Korr DIC objective with silicone oil (Zeiss). The images were cropped to zoom in on the arterial regions using Fiji ImageJ(97) (v.2.16.0).

### Statistical analysis

All statistical analyses were done using GraphPad Prism (v.10). For single-factor experiments, the normal distribution of the data and the residuals were checked, and one-way ANOVA followed by Tukeýs, Dunnett’s, or Sidak’s post-hoc tests were used. For two-factor experiments, two-way ANOVA followed by Tukeýs, Dunnett’s, or Sidak’s post-hoc tests was used. The data were plotted using GraphPad Prism (v.10) or ggplot2 in R. The p-values smaller than 0.1 were denoted on the plots. The details for the statistical methods and the experimental replicates are given in the Fig. legends.

### 1D Model of Hybrid Arterial Wall

To simulate the Notch dynamics in smooth muscle cells (SMCs) within a *hybrid artery* — representing signaling under non-periodic boundary conditions replicating *in vivo* conditions—we adapted an existing computational framework, similar to previous studies(13, 98), we assumed that the first cell in the linear arrangement is an endothelial cell that initiates the signaling cascade through constant emission of Jagged1 (Fig. 4B). SMCs interact with each other as in the previous model, where trans interactions occur between proteins located on the membranes of adjacent cells. Notch and Jagged are both assumed to be evenly distributed across the cell membrane, with half of their total quantity available for binding with each neighboring cell.

Considering the significance of Notch in coronary artery dynamics, we selected the geometrical parameters for a coronary artery as the “hybrid artery” to be simulated, assuming an average wall thickness of 0.2 mm(99). The simulated cell line consisted of 20 cells, assuming a cell size of 0.01mm, consistent with the arterial thickness. The endothelial cell’s constant Jagged1 emission rate was set to 4000 molecules/ℎ, in agreement with previous studies(13, 98).

## Supporting Information Text

### Supporting Methods

#### Cell Culture and Maintenance

Human aortic smooth muscle cells (SMCs) were purchased from Lonza (Cat. #CC-2571) and American Type Culture Collection (ATCC Cat. #PCS-100-012) in passages 3 or 2 (See Table S4 for details). The cells were grown in Smooth Muscle Basal Medium containing SmGM^®^-2 SingleQuots supplements (Lonza Cat. #CC-3182) until passage 4-6 and used within 15 population doublings. Human umbilical vein endothelial cells (HUVECs) were bought from PromoCell (Cat. #C-12200) at passage 1 and grown in Endothelial Cell Growth Medium 2 (PromoCell Cat.#C-22011) until passage number 4. Human embryonic kidney cell line HEK 293T was grown in Dulbeccós Modified Eaglés Medium (Merck Cat. #D6429) supplemented with 10% fetal bovine serum (Gibco Cat. #17914671), 2 mM L-glutamine (Merck Cat. #G7513) and 100 units/ml penicillin/streptomycin (Merck Cat. #P0781). All cells were maintained at 37°C with 5% CO_2_ and kept under 80% confluency at all times unless the experimental setting required otherwise.

#### Ligand Coating

For the Notch activation assays, the recombinant human Jagged1-Fc chimeric ligands (Bio-techne Cat. #1277-JG) were immobilized on the culture plates at 10nM concentration as described previously(1). Briefly, the culture plates were first treated with 50μg/ml protein G (Thermo Fisher Scientific Cat. #21193) in phosphate-buffered saline (PBS) (Biowest Cat. #L0616) overnight at RT. The next day, the plates were washed three times with PBS and blocked with 1% bovine serum albumin (BSA) (Merck Cat. #A9418) in PBS for 1 hour at RT. Then, the plates were treated with the ligands in PBS with 0.1% BSA overnight at cold or 2 hours at RT. The plates were washed three times with PBS before use.

#### RNA interference and Plasmids

For siRNA transfections, 10^4^ SMCs per cm^2^ were plated in 6cm culture dishes and grown overnight. The next morning, the cells were transfected with 25nM Dharmacon ON-TARGETplus SMARTPool siRNAs (Horizon Discovery) for the indicated genes or with the same concentration of non-targeting siRNA (Eurofins Genomics) using RNAiMAX lipofectamine (Thermo Fisher Scientific Cat. #13778150). Briefly, the siRNAs and the lipofectamine solutions were separately prepared in Opti-MEM (Gibco Cat. #31985062). The lipofectamine solution was dropwise added to the siRNA solution and mixed gently by flicking the tubes. The solution was then briefly centrifuged and left to incubate for 15 min at RT. The cells were supplied with fresh growth media. The siRNA mix was dropwise added to the culture plate and swirled gently to mix. The cells were then incubated for 24 or 48 hours according to the experimental setting before use. 10 For plasmid transfection, HEK 293T cells were grown to 70% confluence in 10 cm culture dishes. The indicated plasmids were transfected using jetOPTIMUS transfection kit (Sartorius Cat. #101000006) at 1μg/ml concentration, according to the supplieŕs recommendations. Briefly, the plasmids and the transfection reagent were added to the supplied buffer and mixed using a vortex for 1 second. The tube was centrifuged for 10 s and left to incubate for 10 min. The solution was then dropwise added to the culture plate that was supplied with fresh, warm media. The cells were then incubated overnight before use.

#### Fixation and mounting for microscopy

The cells were washed three times with HBS, fixed with 4% PFA in PBS for 10 min at RT, and mounted using Vectashield with DAPI (4’,6-diamidino-2-phenylindole, dihydrochloride) (Vector Laboratories Cat. #H-1000). The samples were imaged using the 3i CSU-W1 spinning disc confocal microscope.

#### Flow cytometry

The cells were washed three times with HBS and detached using ReagentPack™ subculture reagents (Lonza Cat. #CC50-34) at RT and immediately taken on ice and collected into 1.5 ml centrifuge tubes. The cells were washed with ice-cold HBS three times by centrifuging for 3 min at 500 RCF, 4°C, and resuspending in fresh buffer. The cells were analyzed with BD LSR Fortessa flow cytometer using 488 nm excitation wavelength. The data were analyzed using the FlowJo software (v.10) and presented as mean fluorescence intensity.

#### Spheroid formation

8x10^3^ SMCs were grown in Nuncleon™ Sphera™ low adhesion plates (Thermo Fisher Scientific Cat. #174929) for 72 hours in SMGM-2 media to form mono-culture SMC spheroids. For the co-culture spheroids, SMCs were first grown for 48 hours alone and then for 24 hours with 3x10^3^ HUVECs. The spheroids were washed once with PBS, then fixed and permeabilized in PBS containing 4% paraformaldehyde and 1% Triton X-100 (VWR, Cat. #M143) overnight at cold. They were washed three times for 15 min each in PBS with 0.1% Triton X-100 and incubated with 1:100 mouse CD31 (Thermo Fisher Scientific Cat. #37-0700) and 1:100 rabbit Jagged1 (Cell Signaling Cat. #2620) primary antibodies overnight at cold. The spheroids were washed three times and incubated with 1:400 dilutions of the anti-rabbit Alexa fluor 488 (Thermo Fisher Scientific Cat. #A-11034) and anti-mouse Alexa fluor 555 antibodies (Thermo Fisher Scientific Cat. #A-21422). After three more washes, the spheroids were mounted in glycerol (Merck Cat. #G5516). The samples were imaged using the 3i CSU-W1 spinning disc confocal microscope. The mean Jagged1 signal in confocal slices from the middle of the spheroids was quantified using Fiji ImageJ and normalized by subtracting the intensity of the control samples without primary antibodies. At least four spheroids were analyzed per group for each independent experiment. The normalized signal was presented as a fold change.

#### RNA isolation, cDNA synthesis, and qPCR

RNA was isolated from SMCs using NucleoSpin RNA (Macharey-Nagel Cat. #740955) according to the supplieŕs protocol. The RNA concentration and quality were checked using Nanodrop ND-2000, Thermore Fisher Scientific. The resulting RNA was used for cDNA library preparation using the SensiFAST™ cDNA synthesis kit (Bioline Cat. #BIO-65054). The quantitative polymerase chain reaction (qPCR) using 5x HOT FIREPol EvaGreen qPCR Mix Plus reagent (Labnet Cat. #08-24-00001-10) according to the supplieŕs protocol using Quanstudio Real-Time PCR system, Thermo Fisher Scientific. The qPCR primers for the indicated genes were ordered from TAG Copenhagen (See the Table S2 for the primer sequences). For all primer pairs, a cDNA dilution series, prepared from a cocktail of samples from the same experiment, was included in the same qPCR run to calculate a standard curve. The standard curve was used to calculate the relative mRNA expression.

#### RNA Sequencing and Data Processing

For each sample, a starting quantity of 100 ng RNA was processed. Sample quality was ensured using Agilent BioAnalyzer 2100. Sample concentration was measured with Qubit^®^/Quant-IT^®^ Fluorometric Quantitation, Life Science Technologies. The library preparation was done according to the guidelines from Illumina Stranded mRNA Preparation, Illumina (1000000124518) using Illumina Stranded mRNA Preparation, Ligation, Kit (Illumina). The average fragment size was 335 bp. The base calling was done using the bcl2fastq2 software on NovaSeq 6000. Sequencing was done using Illumina NovaSeq 6000 S1 v1.5, with paired-end sequencing and 2 x 50 bp read length.

The read quality was checked with FastQC (v.0.11.8). The reads were aligned against the hg38 human genome (Illumina iGenome: https://support.illumina.com/sequencing/sequencing software/igenome.html (UCSC)) and annotated using the Rsubread package (v.2.6.4) in Bioconductor (v.3.13), R (v.4.1.0). The gene ids were matched to the gene symbols using the org.Hs.eg.db package (v.3.13.0). Normalization was done with edgeR (v.3.34.1) to produce counts per million (CPM) and reads per kilo read per million (RPKM) for each gene. Pairwise comparisons between groups were performed using ROTS (v.1.20.0) on log2(1+CPM).

#### Western blot

All samples were lysed in 3x Laemmli buffer (0.1875 M Tris, 0.21 M SDS, 30% glycerol, bromphenol blue) containing 3% 2-mercaptoethanol (Merck, Cat. #444233) on ice, scraped into 1.5 ml centrifuge tubes, and boiled at 95°C for 10 minutes. The lysates were separated by SDS-PAGE and transferred onto nitrocellulose membranes (Cytiva, Cat. #15220033). The membranes were checked for loading using Revert™ 700 Total Protein Stain (LI-COR Cat. #926-11016), blocked with 1% w/v milk in PBS with 0.1% Tween 20 (VWR Cat. #M147) for 30 min and probed with the antibodies indicated below. The bands were visualized using SuperSignal™ West Picoplus chemiluminescent substrate kit (Thermo Fisher Scientific Cat. #34580) and imaged with iBright FL1000, Thermo Fisher Scientific. The bands were analyzed using the ImageStudio Lite software (v.6.0), LI-COR. Unless stated otherwise, all bands were normalized to total protein stain. The primary antibodies used: Notch2 1:1000 (Cell Signaling Cat. #5732), Notch3 1:1000 (Cell Signaling Cat. #5276), Jagged1 1:1000 (Cell Signaling Cat. #2620), α-SMA 1:1000(Cell Signaling Cat. #19245), Pdgfrβ 1:1000 (Cell Signaling Cat. #3169), β-tubulin 1:1000 (Cell Signaling Cat. #86298), His-Tag 1:1000 (Cell Signaling Cat. #2365), M2-Flag 1:1000 (Merck, Cat. #F1804), smad2/3 1:200 (Santa Cruz Cat. #sc-133098), and phospho-smad2 1:1000 (Cell Signaling Cat. #3108), β-actin 1:10000 (Merck Cat. #A1978). The horse radish peroxidase-conjugated secondary antibodies used: anti-rabbit 1:10000 (Vector Laboratories, #PI-1000), anti-mouse 1:10000 (Vector Laboratories, #PI-2000), Veriblot for IP detection 1:500 (Abcam, Cat. #ab131366).

#### Re-analysis of single-cell RNAseq data

For the vascular cells from different organs, previously published normalized datasets from the Betsholtz lab(2–4) were downloaded from Gene Expression Omnibus under the accession codes GSE98816 (brain), GSE20100 (lung), GSE210087 (heart), GSE210013 (colon), and GSE21000 (aorta). The expression matrices were read using the general Seurat scRNAseq analysis pipeline (v.4.4). For the datasets except the brain data, the ENSEMBL IDs were converted to gene symbols using Org.Mm.eg.db (v.3.20.0) and AnnotationDbi (v.1.68.0), and the duplicates and the entries without symbols were removed. Each dataset was then processed according to the Seurat workflow. Gene expression was normalized to 10000 counts and log-transformed. The clusters were created using the FindClusters function with the resolution set to 0.5 for the aorta dataset and 1 for the rest. The UMAP was calculated for brain, lung, heart, colon, and aorta datasets from the top 20, 25, 20, 20, and top 15 PCs, respectively. The cluster annotations were partly adapted from Muhl et al.(4). SMCs, Acta2, Myh11, Tagln, and Cspg4 were used as SMC markers, while Chrdl1, Hhip, Shisa3, Ccl19, Chrm2, Bdkrb2, and Abcc9 were used to filter out the non-arterial SMCs and pericytes. For the heart dataset, the cluster positive for Itgbl1 was annotated as aortic SMCs proximal to the heart. The clusters positive for Pecam1 and Kdr and negative for Prox1 were annotated as vascular endothelial cells. The clusters were extracted from the count matrices using the annotated cell IDs from the Seurat objects. The SMC and the EC data were processed separately. The datasets were filtered only to include the common genes among the different organs. The extracted matrices were re-normalized individually using “RC” transformation. No data integration or batch correction were deemed necessary as the datasets were used only for the comparison of single genes.

The processed mesoderm dataset was downloaded from Zebrahub. The expression matrix and the metadata files were extracted using the Scanpy tool in Python and used to create a Seurat object. This dataset was already log-normalized and scaled and did not contain the raw counts. Hence, the PCA analysis was done manually using the first 30 components. The clusters were found using the resolution set to 0.2, and the UMAPs were found using the top 15 PCs. Datasets from different timepoints (batches) were integrated using harmony package. Markers for all clusters were found using the FindAllMarkers command. The markers for non-vascular associated SMC markers and cycling cells were checked using the Daniocell(5) single-cell RNAseq database. Based on this, the clusters highly expressing, desmb, nkx2.3, rem1, and tlx1 were used to filter out the non-vascular SMCs (Figure S4A). The remaining SMC clusters were annotated based on smooth muscle marker and notch3 expression. The VlnPlot function was used to plot the normalized counts based on the new annotations.

#### Computational Model for Extrinsic Ligand Induction

To simulate the experimental scenarios of Notch signaling in a cluster of SMCs, we extended a previous computational framework of Notch signaling (AB model developed by Loerakker et al.(6) and Ristori et al.(7), based on Sprinzak et al.(8) and Boareto et al.(9)) in a periodic one-dimensional array of SMCs, by incorporating the separate effects of Notch2-Jagged1 and Notch3-Jagged1 binding on the signaling dynamics. The levels of key proteins (Notch2, Notch3 Jagged, NICD) were predicted for each cell, with interactions allowed between neighboring cells on either side. The temporal changes in Notch (N), Jagged (J), and NICD (I) concentrations within cell i at time t were modeled using the following differential equations:

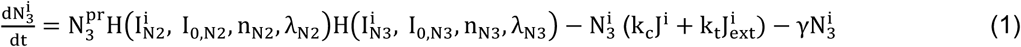

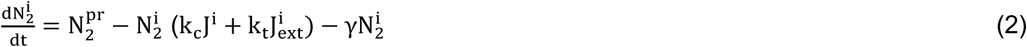

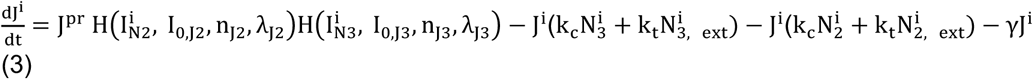

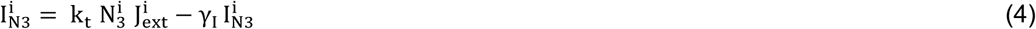

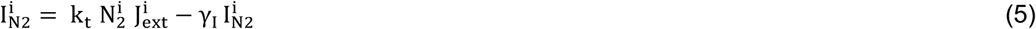

Here, N_2_, N_3_, J represent the Notch2, Notch3, and Jagged1 protein levels, and I_N3_, I_N3_ represent the NICD content of the cell. 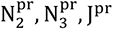 are the basal production rates of Notch2, Notch3 and Jagged1, respectively. k_c_, k_t_ denote the cis-inhibition and trans-activation strength parameters, respectively. γ represents the degradation rate of Notch and Jagged and γ_I_ is the degradation rate of NICD. The effects of transactivation on the production rate of Notch3 and Jagged1 from NICD2 and NICD3 were incorporated with shifted Hill functions in the following form:

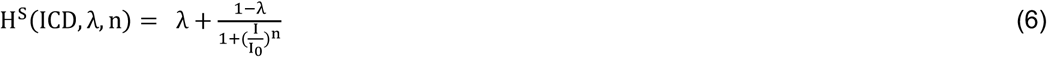

with I denoting the NICD content (NICD_N2_ or NICD_N3_) and λ defining the changes in protein production in response to Notch2/Notch3 transactivation. The parameter I_0_ defines the transition point of the Hill function from convex to concave and the parameter n determines the sensitivity of protein production to the NICD content. A schematic illustration of the interactions in the model is shown in Figure S3B.

The initial values for the protein content of Notch2, Notch3, Jagged, and NICD for each cell were randomly generated between 0 and 6000 molecules, similar to Loerakker et al.(6) and Ristori et al.(7). The trans interactions occur with proteins located on the membranes of neighboring cells. Notch and Jagged were assumed to be distributed equally over the cell membrane, where half of the amount is available for binding with each of its neighbors. To replicate the experimental environment of a large number of closely packed cells in a cluster, periodic boundary conditions were applied to the first and last cells in the simulated array. Hence the first cell was treated as a neighbor to the last cell, creating a continuous loop, thereby eliminating artificial edge effects that would otherwise arise if the array had distinct boundaries (Figure S3C). For all other cells, the external Notch and Jagged were defined in the following form:

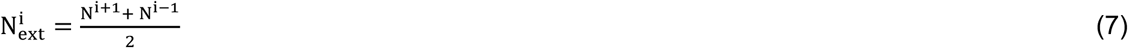

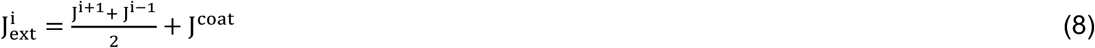

Where 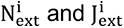represent the external Notch[2/3] and Jagged1 for cell i that is available to for Jagged-Notch signaling, respectively, and J^coat^ is the constant external Jagged coating.

In line with the experimental conditions, we simulated three different protein manipulation setups: (1) Control, with both Notch2 and Notch3 production; (2) siN2, with Notch2 production knocked down; and (3) siN3, with Notch3 production knocked down. These three scenarios were also tested with external Jagged1 coating, resulting in three additional setups: Control_J, siN2_J, and siN3_J, giving a total of six experimental scenarios. Each simulation was repeated 25 times, and all simulations reached a steady-state solution after a simulated time of 15 hours. The stable protein content values in the periodic model were found to be insensitive to the to the time increment when chosen ≤0.01 of the explicit time integration scheme and the number of cells when chosen ≥10 cells, hence a time increment of 0.01 and 10 cells were adopted. The final protein production levels were averaged across cells and repeated simulations for each experimental scenario.

Most of the parameter values regulating the dynamics of Jagged1–Notch2/3 signaling in SMCs were adopted from Boareto et al.(9). The basal production rates of Notch2 and Jagged1 were assumed to be half that of Notch3, which was set at 1000 h^−1^, since based on RNAseq data, SMCs expressed 2-fold more copies of NOTCH3 than NOTCH2 at 24h (Figure S3 A). In addition, they seemed to express almost no other ligand than JAG1. The strengths of trans-activation (k_t_) and cis-inhibition (k_c_) were assumed 5 × 10^−5^ and 5 × 10^−4^, respectively(6, 9). Hill function parameters were selectively optimized, with only the transition parameters I0_J3_ and I0_N3_ and sensitivity parameters n_N2_ and n_N3_ adopted from previous models(6, 9). The remaining parameters of the shifted Hill function were optimized to account for their potential influence on the results (see Table 1).

**Table S1.**
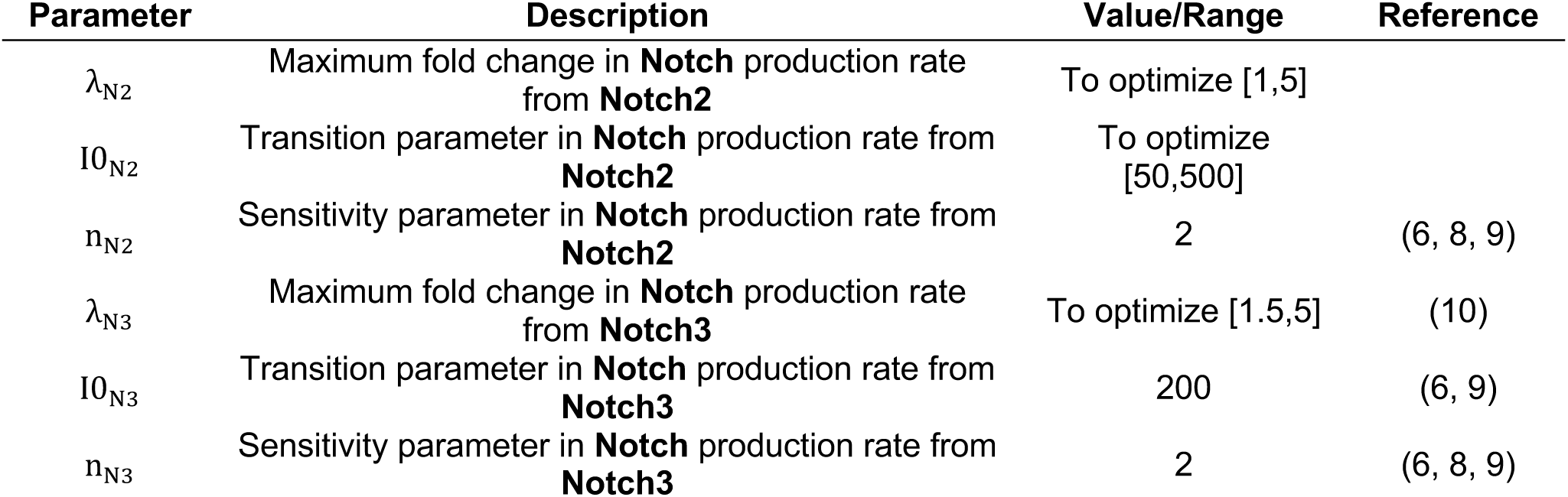
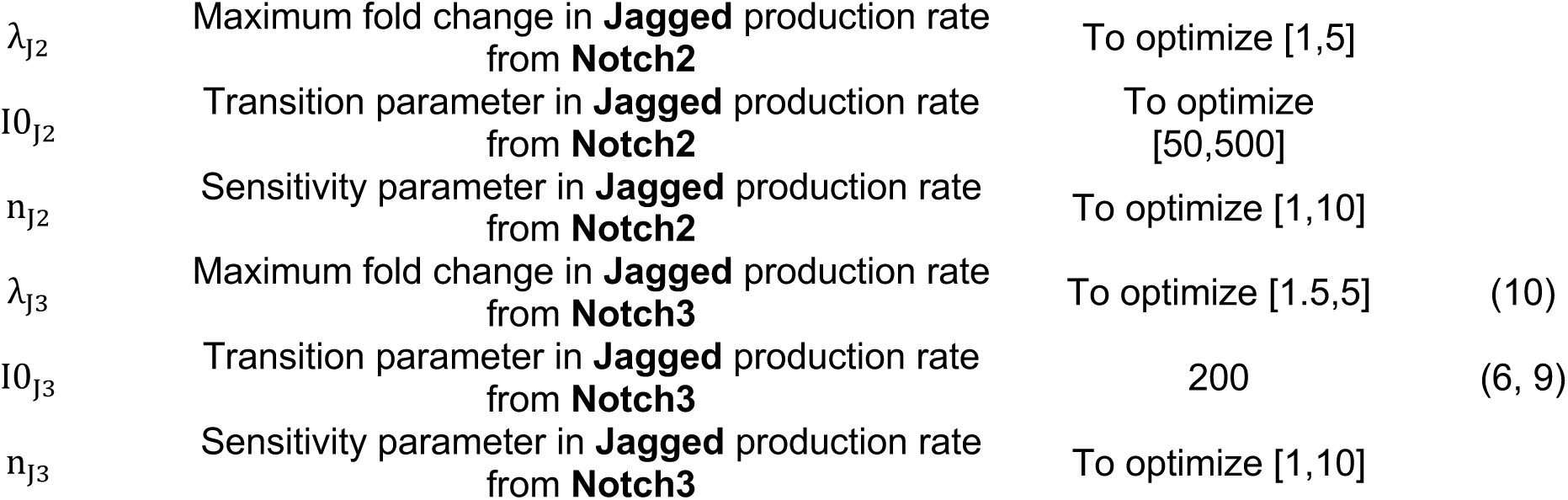
Parameters of the shifted hill function characterizing the upregulation of Notch3 and Jagged 1 by NICD2 and NICD3 in the computational model.

We employed a Genetic Algorithm (GA) to optimize a set of eight parameters (λ_N2_, I0_N2_, λ_N3_, I0_N3_, n_N3_, λ_J2_, I0_J2_, n_J2_, λ_J3_, I0_J3_, n_J3_) in the computational model of Notch signaling, aiming to minimize the Mean Squared Error (MSE) between simulated and experimental qPCR data at 24h from Figure 2 G for transmembrane proteins. The simulations were performed using MATLAB (The MathWorks Inc., v.R2025a). The objective function was defined to calculate the MSE between the normalized simulated outputs (averaged for cells at 24h) and the experimental data, using the Notch2-Notch3 signaling model. The GA was run with a maximum of 100 generations and the stopping criteria were set to three consecutive generations with no improvement, and a tolerance of 10^−4^. The objective function calculated the Mean Squared Error (MSE) between simulated and experimental data for Notch2 (N2), Notch3 (N3) and Jagged1 (J1) relative expressions. As Notch2 is not influenced by Hill function parameters (Eq 2), the MSE was computed by summing the squared differences normalized values 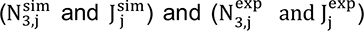, where j denotes the index of the experimental scenario. Thus, the total MSE serves as the objective function (CF) that quantifies the difference between the two datasets.

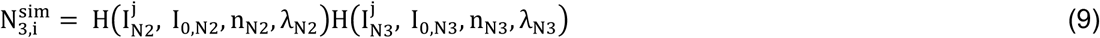

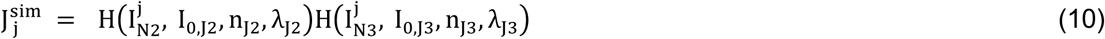

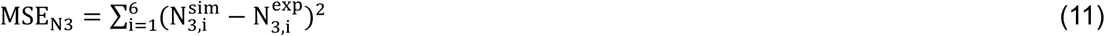

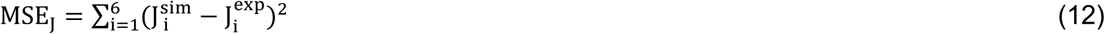

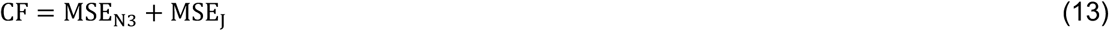

The optimization results including the best parameter values, number of iterations and the minimum cost value are given in Table 2.

**Table S2.** *Optimized parameters of the Notch signaling model using the Genetic Algorithm under constraints of* Table 1.

| Optimized Parameters |  |  |  |  |  |  |  | Iterations | Cost |
| --- | --- | --- | --- | --- | --- | --- | --- | --- | --- |
| $\lambda_{N2}$ | $I0_{N2}$ | $\lambda_{N3}$ | $\lambda_{J2}$ | $I0_{J2}$ | $n_{J2}$ | $\lambda_{J3}$ | $n_{J3}$ | | |
| **2.57** | 50.24 | 1.56 | 4.52 | 76.38 | 1.89 | 1.82 | 1.76 | 65 | 0.68 |

The optimization results showed a stronger influence of NICD2 on transmembrane protein production (Notch and Jagged), compared to that of NICD3 (λ_J2_, λ_N2_ > λ_J3_, λ_N3_). Additionally, I0_J2_, I0_N2_ < I0_J3_, I0_N3_ = 200, which shows that NICD2 induces Notch and Jagged at lower thresholds than those of NICD3. To verify these findings, we repeated the optimization procedure with constraints that enforced the opposite conditions:

(1) NICD3 was set to have a stronger influence on the production of Jagged1 and Notch3 than NICD2 (λ_J3_, λ_N3_ > λ_J2_, λ_N2_)
(2) NICD3 was assumed to induce Notch3 and Jagged1 production at lower thresholds than NICD2 (I0_J3_, I0_N3_ > I0_J2_, I0_N2_)

The results of the verification run are shown in Table 3, where both constraints led to significantly higher minimum costs in the optimization algorithm, indicating that these alternative scenarios deviated from the experimental mean protein levels (see Table 3 and Figure S3G).

**Table S3.**
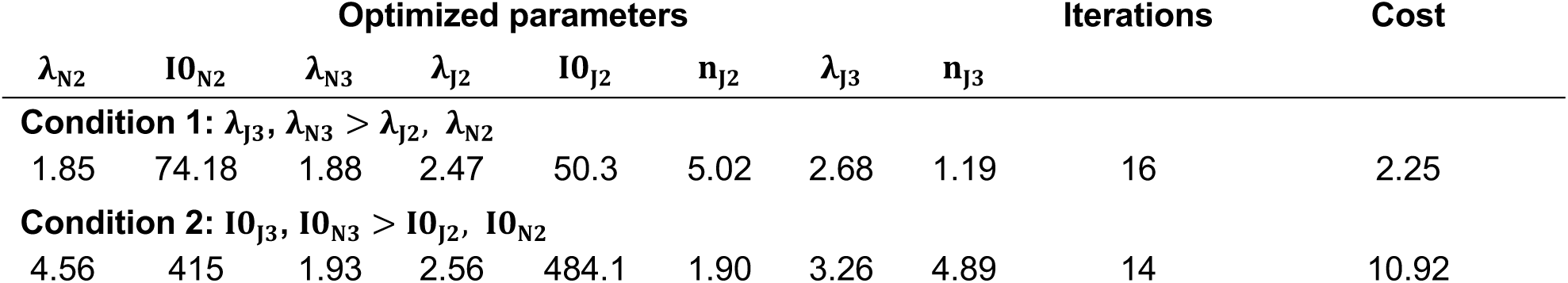
Optimized parameters of the Notch signaling model after enforcing opposite conditions of the original results from Table 2.

**Fig. S1.**
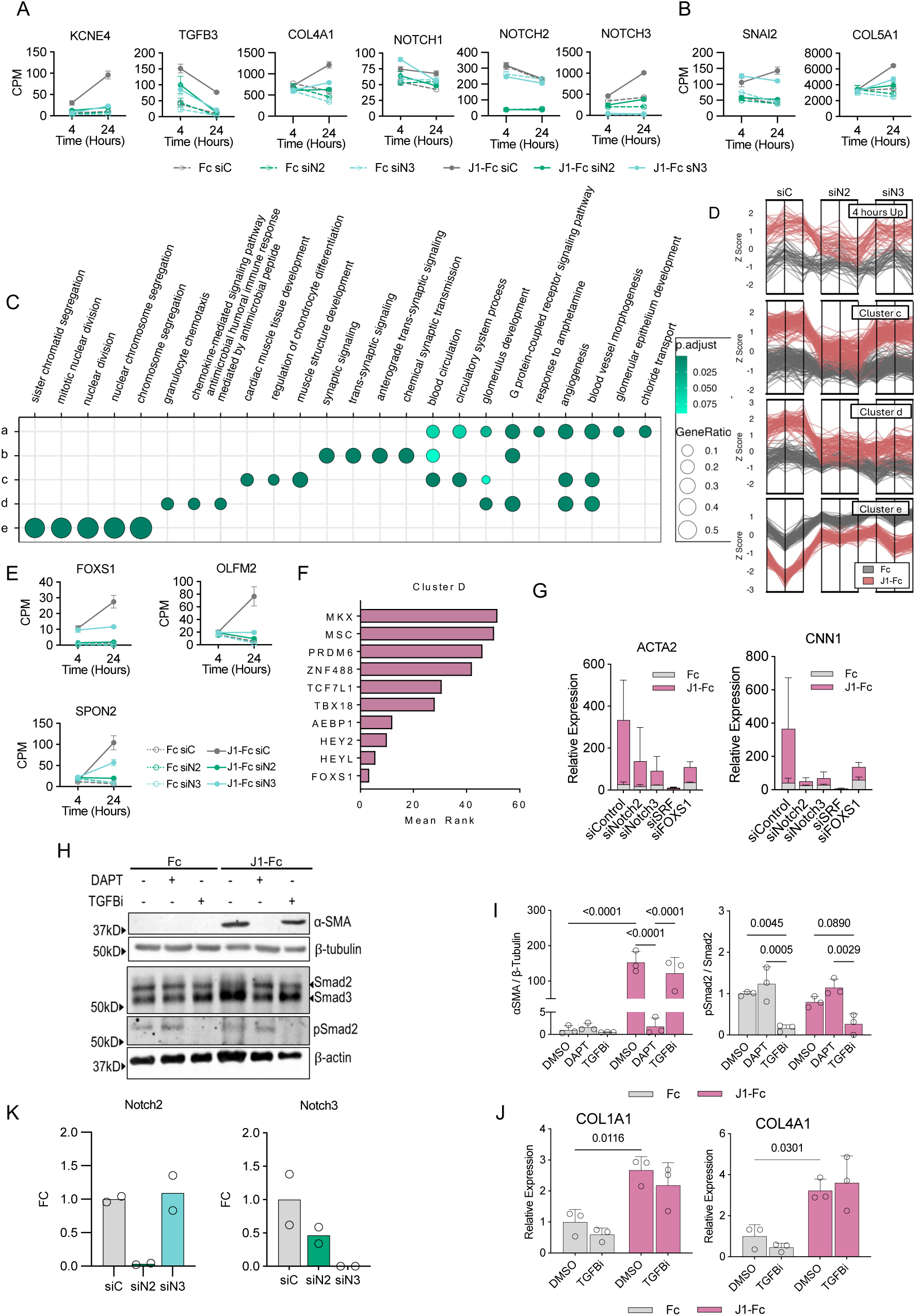
Supplemental data relating to the bulk RNAseq and J1-induced gene expression. (Related to Figure 1 and 2) (A, B) Selected genes from cluster d (A) and cluster c (B), regulated by Jagged1-Notch2/Notch3 signaling. NOTCH1, NOTCH2, and NOTCH3 mRNA levels are given for reference. (C) Gene ontology analysis of the clusters from 24h as listed by the “clusterprofiler” package. (D) Behavior of the gene clusters upon different treatments: genes upregulated on immobilized J1-Fc at 4h, 24h clusters c, d, and e, respectively. (E) Expression of FOXS1, OLFM2, and SPON2 based on RNAseq. FOXS1 was one of the top 10 upregulated genes by J1-Fc stimuli at both time points (please see also Figure 1 C) (mean+SD, n=3 independent experiments). (F) Genes from RNAseq cluster d were analyzed for transcription factor co-expression using the ChEA3 web tool. The top 10 possible upstream transcription factors as listed by ChEA3 are listed with their mean rank (the lower the better) in the bar chart. (G) SMCs were treated with the indicated siRNAs for 48h and grown on immobilized J1-Fc for another 48h. The indicated mRNAs were analyzed using qPCR, normalized to UBC, and presented as relative expression (mean+SD, n=2 independent experiments, one-way ANOVA followed by Dunnett’s test for multiple comparisons to the J1-Fc siControl. The comparisons of J1-Fc-treated samples with p values smaller than 0.1 are shown on the corresponding bars). (H-J) SMCs were grown on immobilized J1-Fc (72h for WB and 24h for qPCR) with 10μM DAPT, TGFβi, or DMSO and analyzed for the indicated molecules. The αSMA and phopho-Smad2 signals were normalized to β-tubulin, and total Smad2, respectively, and presented as fold change (I). For qPCR, the relative expression was normalized to UBC and presented as fold change (J) (mean+SD, n=3 independent experiments, one-way ANOVA followed by Šídák’s test for multiple comparisons. Pairwise comparisons with p values smaller than 0.1 are shown on the plots.).

**Fig. S2.**
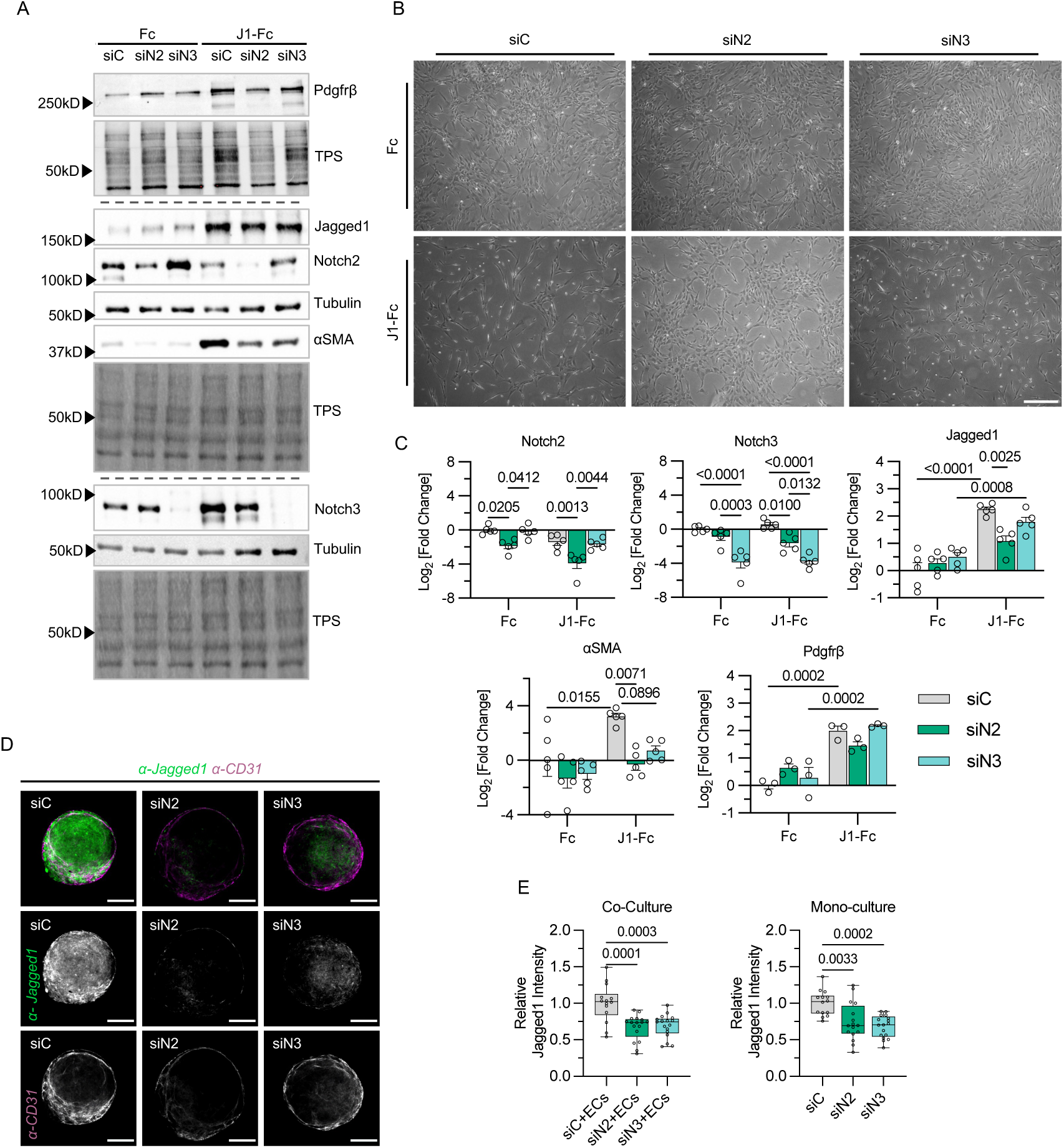
SMC differentiation and Notch activation assays. (Related to Figure 1 and 2) (A-C) SMCs were treated with the indicated siRNAs for 48h, then grown on immobilized J1-Fc for 72h and imaged (C) before immunoblotting for the indicated proteins(A). The signal was normalized to protein loading and presented as log2 fold change (B) (mean+SEM, n=5 (n=3 for Pdgfrβ) independent experiments, two-way ANOVA followed by Tukeýs test for multiple comparisons. Pairwise comparisons with p values lower than 0.1 are indicated on the plots). (D, E) Notch2KD or Notch3KD SMCs were grown as spheroids in low-adhesion U-bottom plates for 72h (monoculture) or 48h before another 24h with human umbilical vein endothelial cells (co-culture). The spheroids were fixed and stained for Jagged1 or CD31. The representative confocal slices (D) and their mean Jagged1 intensity are shown (E) (Scale bars=150μm, error bars represent the min-max range, n=3 independent experiments with at least 4 spheroids per condition, one-way ANOVA followed by Tukeýs test for multiple comparisons. Pairwise comparisons with p values lower than 0.1 are indicated on the plots.).

**Fig. S3.**
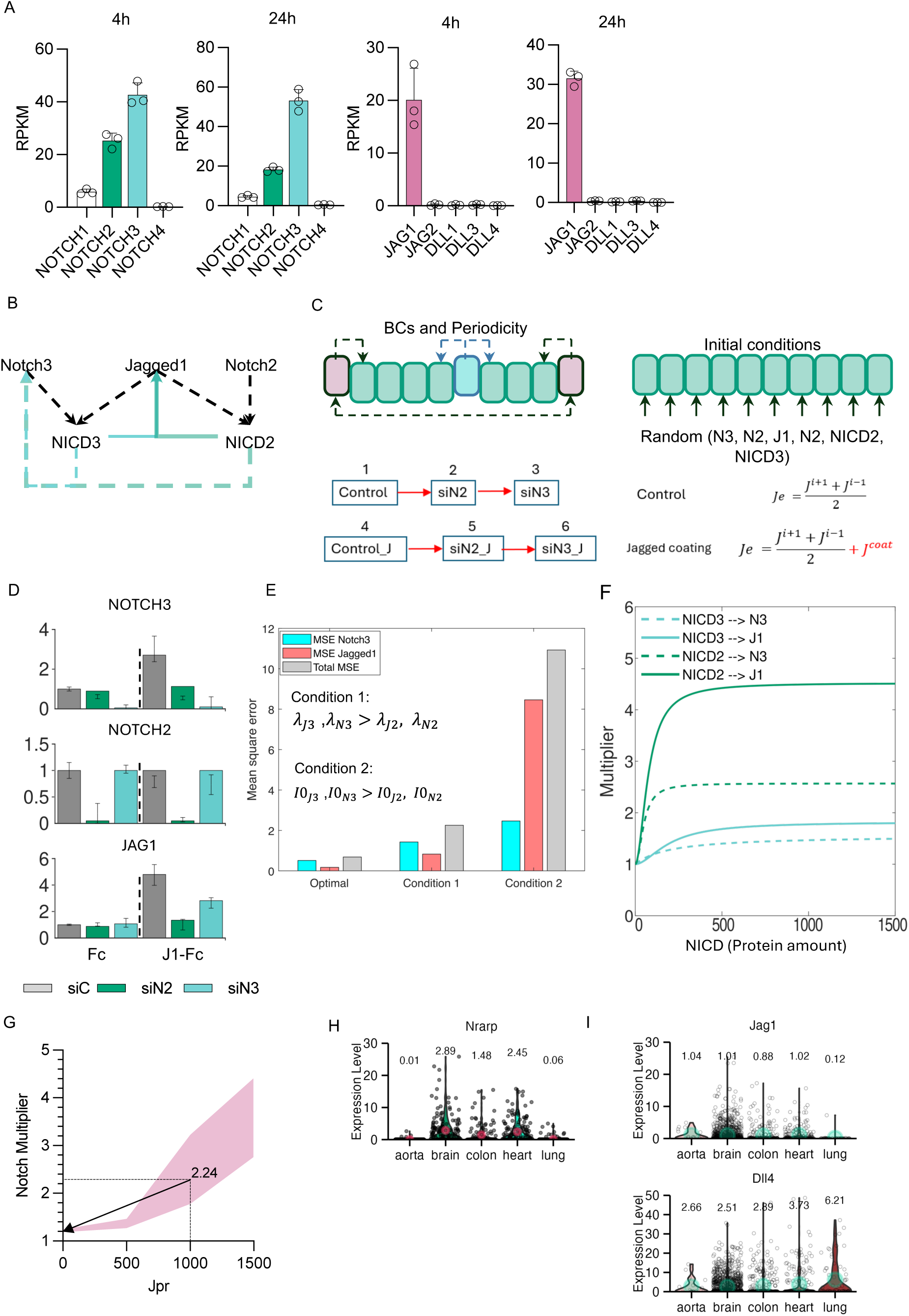
Supplemental data and illustrations relating to the computational model. (Related to Figure 3 and 4) (A) Basal expression of different Notch receptors and ligands in SMCs at 4h and 24h siC- and Fc-treated SMCs based on RNAseq (mean+SD). (B) Diagram depicting the causative relationship between Notch2, Notch3, and Jagged1 included in the theoretical framework used for building the computational model. Line weight denotes the strength of the interaction as informed by the experimental data from Figure 2. (C) Schematic one-dimensional array of vascular SMCs with periodic boundary conditions and initial random values of transmembrane and intracellular proteins (top). Experimental scenarios of protein manipulation with and without jagged coating with external jagged derivation for the cell (bottom). (D) Model validation was performed by enforcing assumptions (conditions 1 and 2) opposite to those the experimental data suggests (optimal). The simulated means against the experimental data (error bars) and the mean squared error for each condition are presented as bar plots. (E) Simulated results from the model showing the levels of the Notch signaling components in response to Jagged1 stimuli following parameter optimization. Bars and error bars represent simulated fold changes and the standard error in fold changes from the experimental data (24h) from Figure 2 G. (F) Hill curves resulting from the simulation with optimized parameters, showing the causative effect of Notch2 or Notch3 ICD on the production of Jagged1 and Notch3 as Notch signaling targets. (G) The predicted Notch signaling multiplier range in the aortic SMCs, was calculated using the ranges N2pr(428:642)(1/h) and N3Pr(857:1071)(1/h). Using N2Pr=500, N3Pr=1000, and Jpr=1000 produces a multiplier of 2.24. Jpr=0 reduces the multiplier to approximately 1. (H, I) Normalized expression of Notch target gene Nrarp in arterial SMCs (H) and Notch ligands Jag1 and Dll4 in vascular endothelial cells (I) of different mouse organs extracted from scRNAseq data (Muhl et al.; aorta, heart, colon, and lung) and (Vanlandewijck et al.; brain). The circles and the text above the violin plots denote the mean expression.

**Fig. S4.**
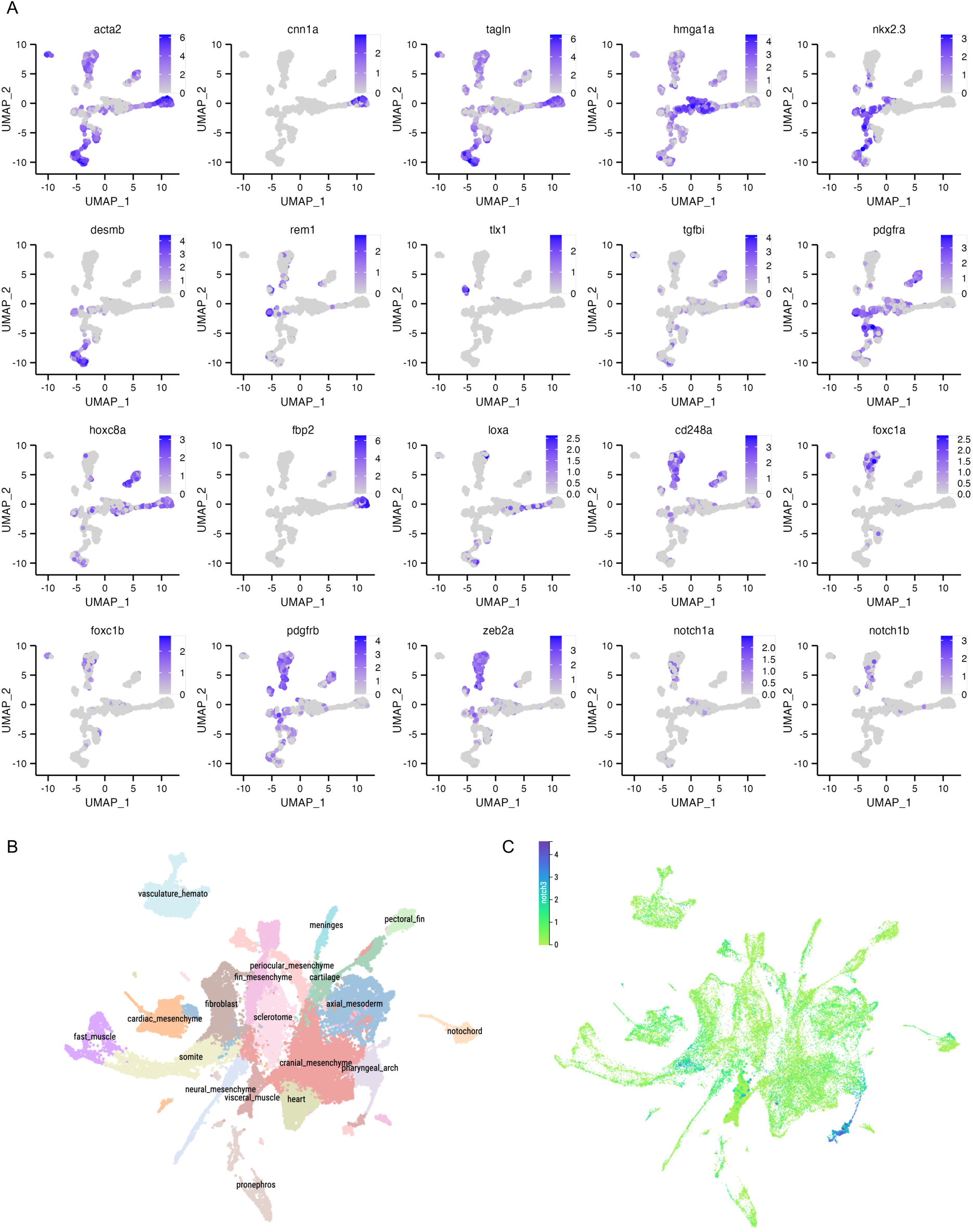
Feature plots of selected genes related to. Fig. 5 **A-F and the clustering of the original mesoderm dataset from Zebrahub.** *(A) Expression levels of smooth muscle markers (acta2, tagln, and cnn1a), cycling SMC marker hmga1a, visceral SMC markers (nkx2.3 and desmb), spleen SMC marker tlx1, fibroblast-like cell markers (tgfbi and pdgfra), positional markers (hoxc8a, hoxc1a, foxc1a-b, fbp2, loxa) and pericytic markers (cd248a, pdgfrb, zeb2a, and abcc9). (B, C) Expression of notch3 across Zebrahub mesoderm clusters – smooth muscle clusters indicated with larger dots*.

**Table S4.**
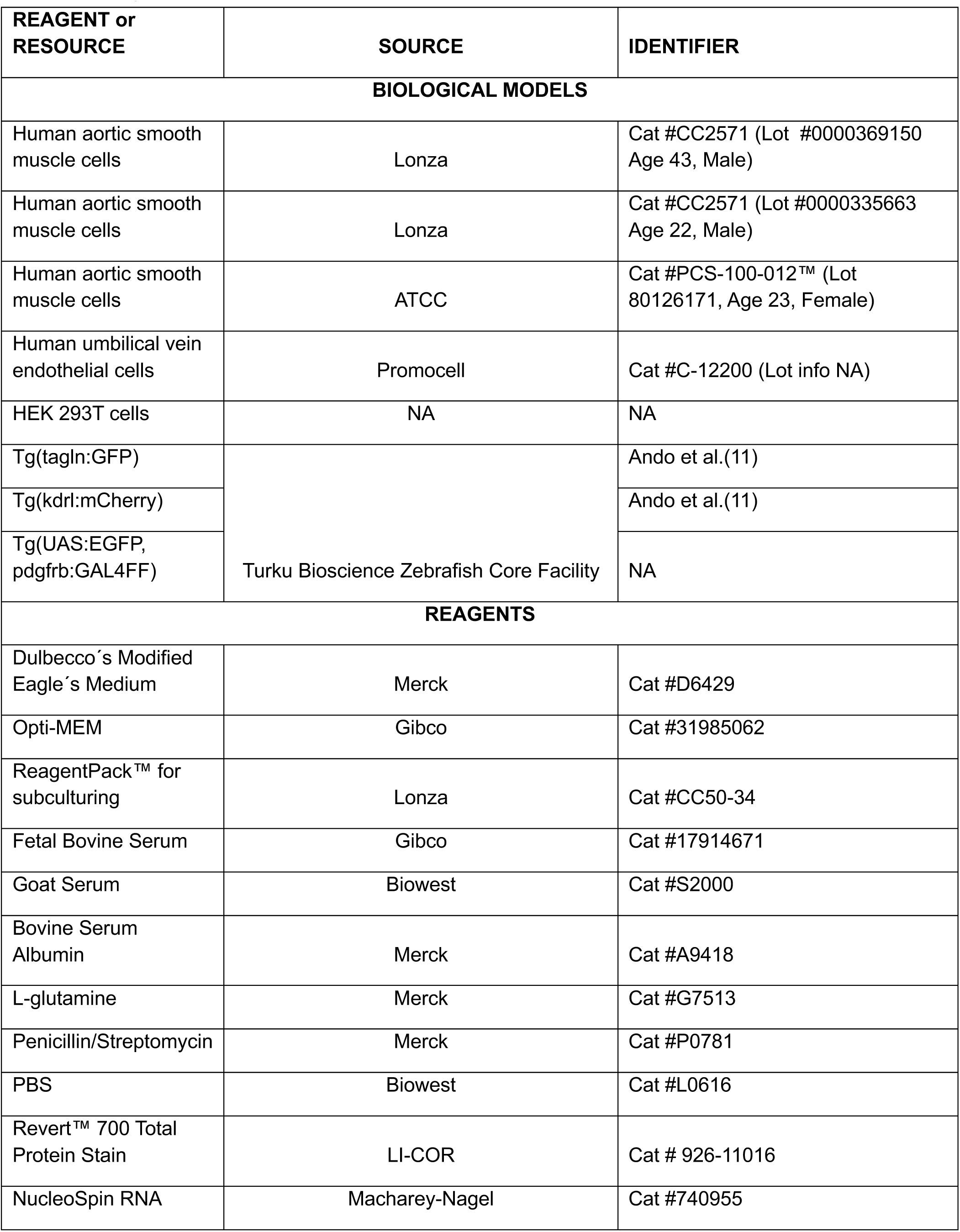
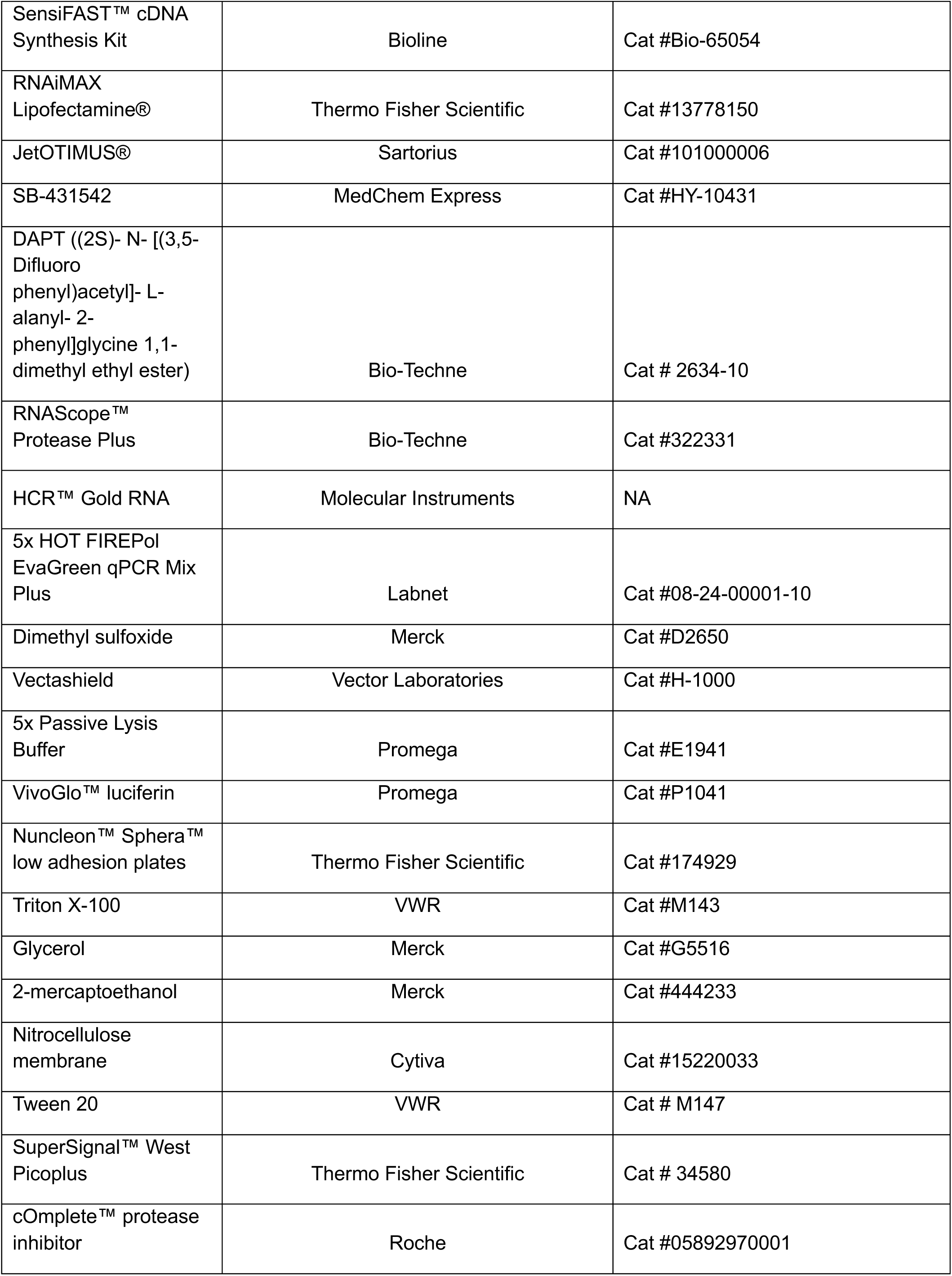
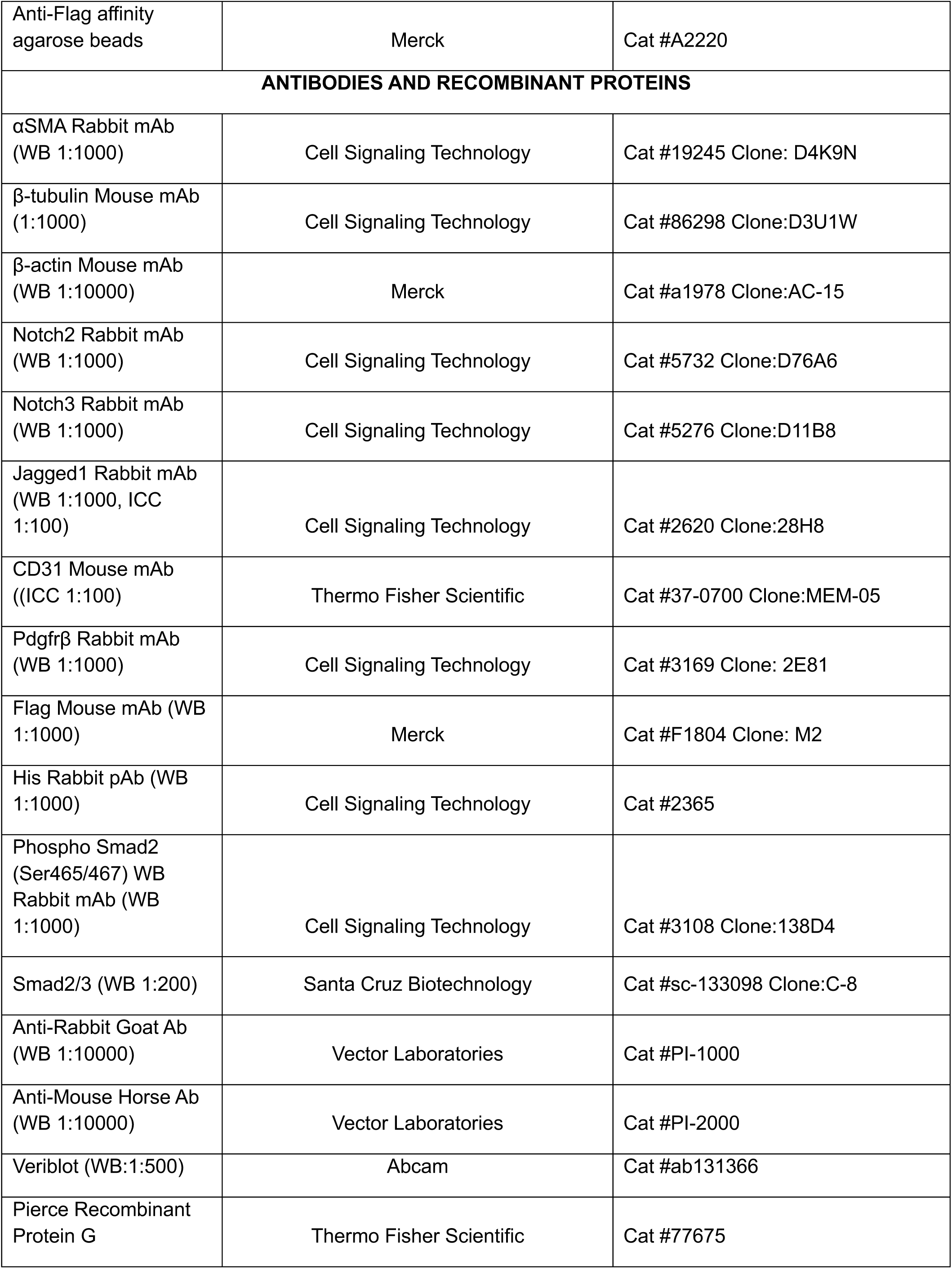
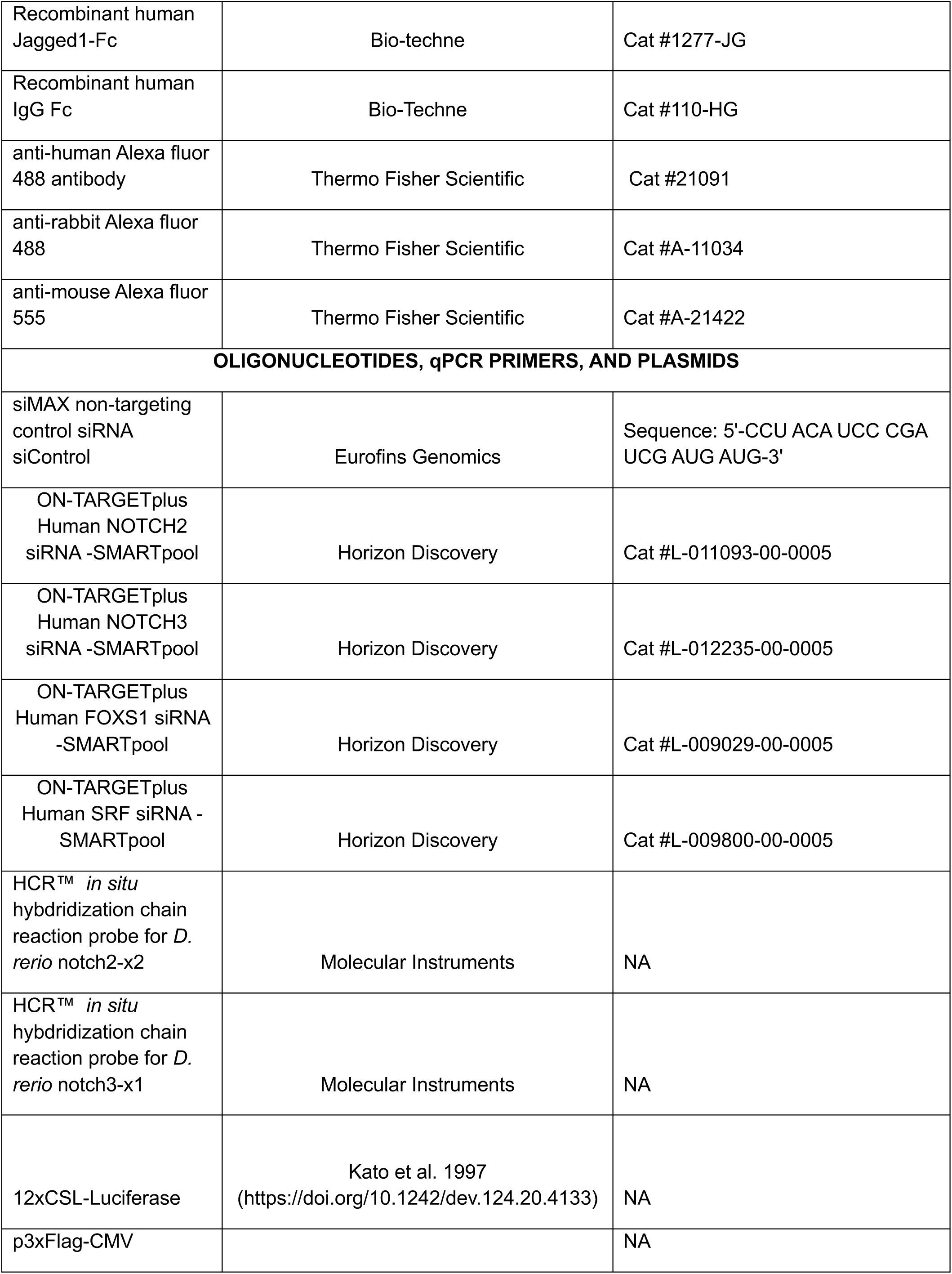
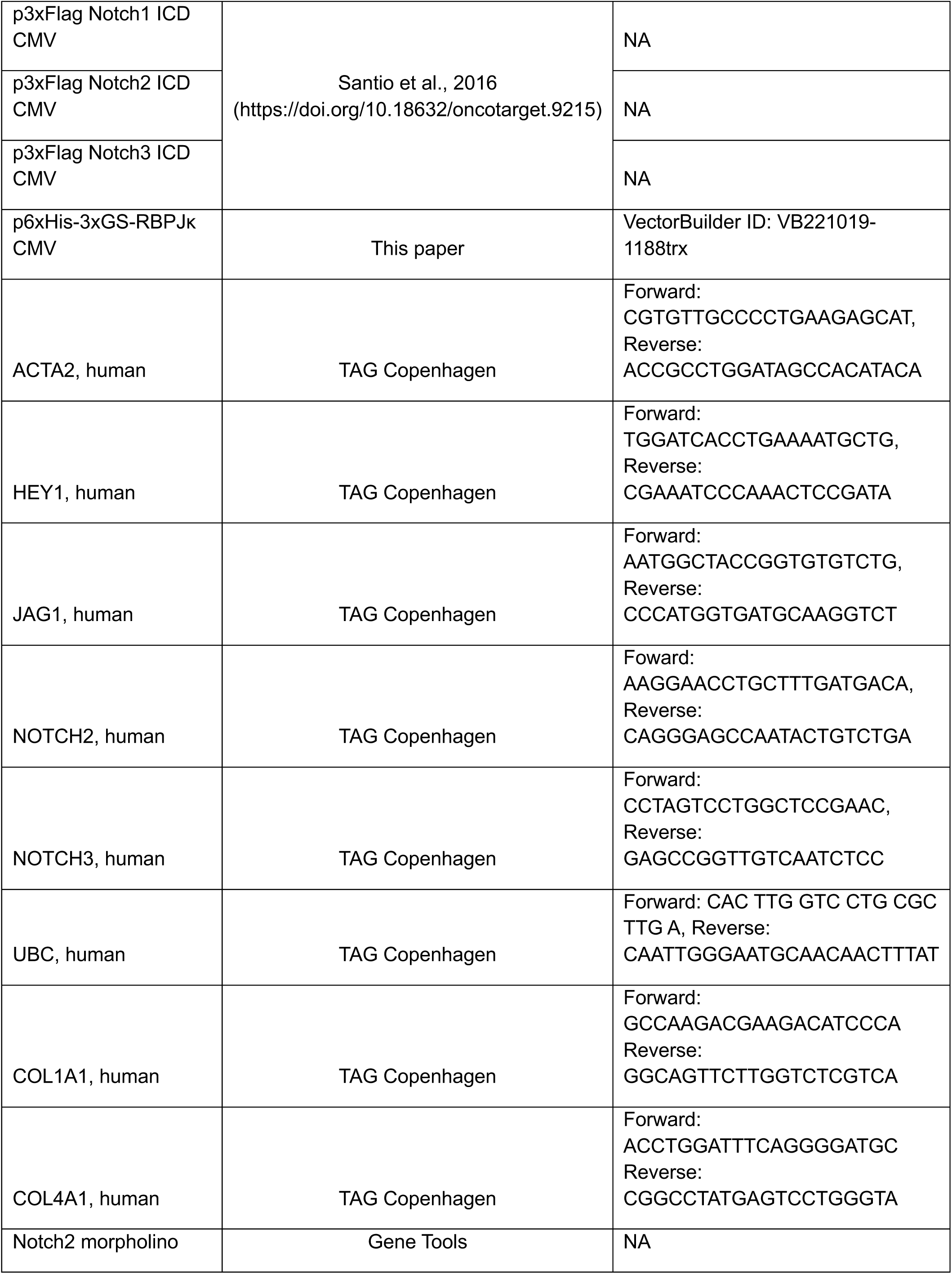
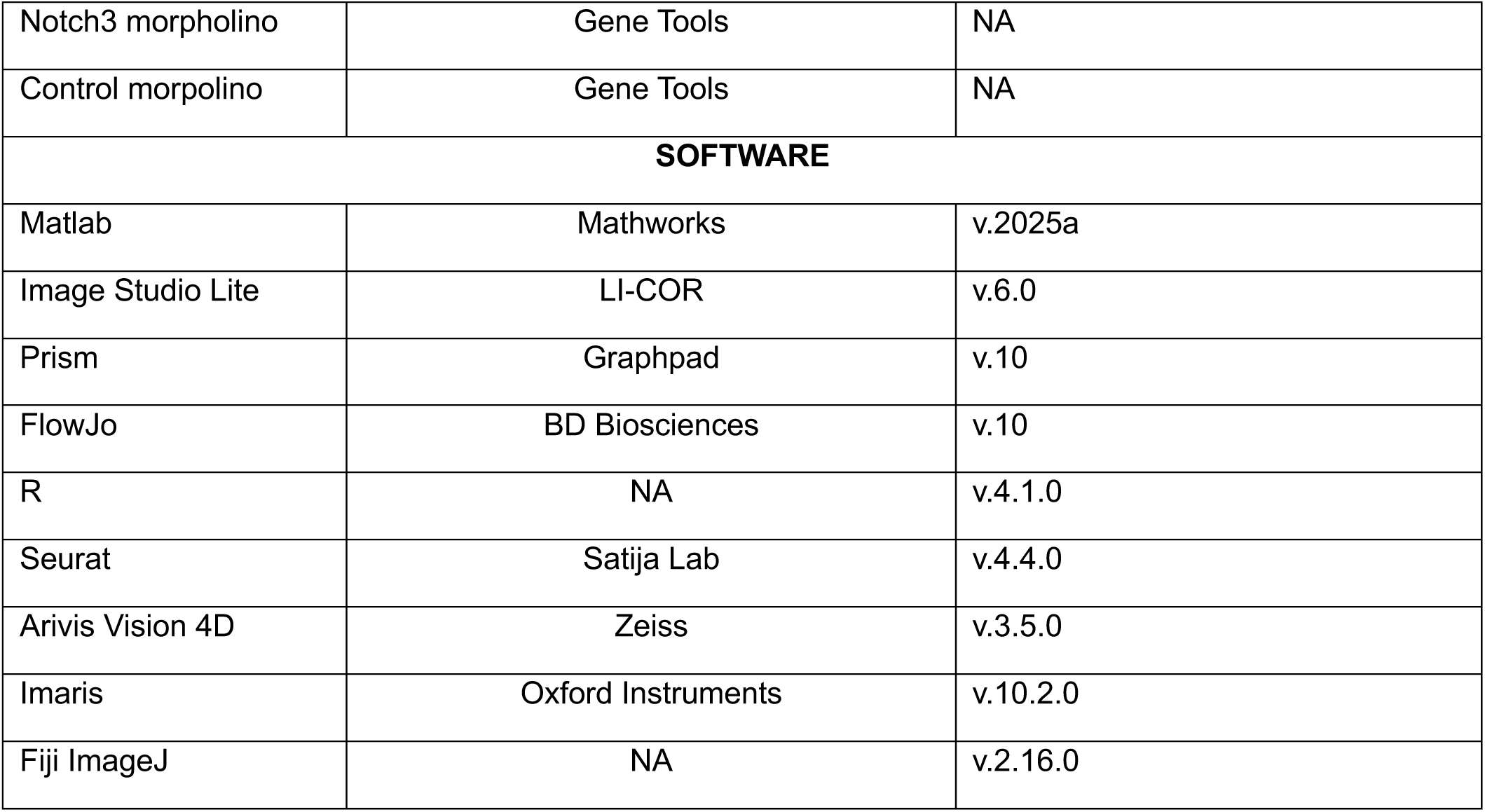
Reagents and other resources

**Dataset S1 (Supplemental_Data_Table_S1.xlsx).** Differentially expressed genes in non-targeting control siRNA-transfected SMCs upon J1-Fc induction and their clustering.

